# Slit-Robo Signalling Establishes a Sphingosine-1-Phosphate Gradient to Polarise Fin Mesenchyme and Establish Fin Morphology

**DOI:** 10.1101/2021.12.07.471519

**Authors:** Harsha Mahabaleshwar, P.V. Asharani, Tricia Loo Yi Jun, Shze Yung Koh, Melissa R. Pitman, Samuel Kwok, Jiajia Ma, Bo Hu, Fang Lin, Xue Li Lok, Stuart M. Pitson, Timothy E. Saunders, Tom J. Carney

**Author notes:** H.M., and A.P.V contributed equally to this work.

## Abstract

Immigration of mesenchymal cells into the growing fin and limb buds drives distal outgrowth, with subsequent tensile forces between these cells essential for fin and limb morphogenesis. Morphogens derived from the apical domain of the fin, orientate limb mesenchyme cell polarity, migration, division and adhesion. The zebrafish mutant *stomp* displays defects in fin morphogenesis including blister formation and associated loss of orientation and adhesion of immigrating fin mesenchyme cells. Positional cloning of *stomp* identified a mutation in the gene encoding the axon guidance ligand, Slit3. We provide evidence that Slit ligands derived from immigrating mesenchyme act via Robo receptors at the Apical Ectodermal Ridge (AER) to promote release of sphingosine-1-phosphate (S1P). S1P subsequently diffuses back to the mesenchyme to promote their polarisation, orientation, positioning and adhesion to the interstitial matrix of the fin fold. We thus demonstrate coordination of the Slit-Robo and S1P signalling pathways in fin fold morphogenesis. Our work introduces a mechanism regulating the orientation, positioning and adhesion of its constituent cells.

## INTRODUCTION

During limb formation, anisotropic growth along the proximal-distal axis results in a flat, paddle-shaped limb bud. How signalling between constituent cells and the biophysical properties of the forming limb are coordinated to attain this morphology has attracted much speculation (Hopyan et al., 2011). Limb bud mesenchyme migration, morphology and adhesion are highly polarised through Apical Ectodermal Ridge (AER) derived signals, including Wnt5a (Gros et al., 2010). This results in filopodial protrusions which orientate radially towards the ectoderm, with a distal bias, and directs polarised orientation, cell division and convergent extension, and thus orientated limb outgrowth (Hopyan *et al*., 2011; Wyngaarden et al., 2010). Furthermore, both tensional forces and a distal-proximal gradient of cell adhesiveness along the limb bud also regulate limb morphogenesis (Lau et al., 2015; Wada, 2011). It is important to understand the processes driving mesodermal cell polarisation, migration and organisation in the limb, and the biophysical properties they impart.

The limb mesenchyme can exert morphogenetic tension on the limb bud extracellular matrix (ECM) through contractility (Martin and Lewis, 1986; Oster et al., 1983). Further, the migration of limb mesenchyme has been proposed to be influenced by haptotactic forces (Oster *et al*., 1983), although this has not been demonstrated *in vivo*. A range of diverse cues alter the adhesive and contractile properties of mesenchymal cells. The soluble phospholipid, sphingosine-1-phosphate (S1P) promotes cell migration, adhesiveness, and myosin based contractile tension in mesenchymal cells and fibroblasts (Hinz, 2016; Hobson et al., 2001; Kanazawa et al., 2010; Wang et al., 1997). S1P signals through G protein-coupled receptors (S1PR1–5), which activate intracellular signalling effectors, including Rho GTPase via the heterotrimeric G-protein Gα_12/13_ (Lee et al., 1998; Wang *et al*., 1997). S1P levels are regulated by dedicated kinases (SPHK1 and SPHK1) or phosphatases (SPP1 and SPP2) (Pitson, 2011), whilst S1P is secreted from source cells by Spinster2 (Spns2) homologues (Osborne et al., 2008). The pathways defining the regulation of intra- and extracellular S1P levels are not fully elucidated.

The importance of S1P in regulating cell behaviour and morphogenesis is demonstrated in zebrafish mutants for *s1pr2*, *spns2*, and *MZsphk2*, which all display cardia bifida, and highlight a role for extracellular S1P in endoderm and cardiac mesoderm migration (Kupperman et al., 2000; Mendelson et al., 2015; Osborne *et al*., 2008). In addition, these mutants all display larval fin blistering, affecting both pectoral and caudal medial fins through an undefined mechanism.

Here, we characterise the zebrafish mutant, *stomp* (*sto*), which shows blisters within the fin folds, similar to those seen in S1P pathway mutants. Surprisingly *sto* corresponded to mutations in the secreted axon guidance protein, Slit3. We show that Slit-Robo signalling is required for S1P potency in the fin fold and that S1P acts to polarise immigrating fin mesenchyme, altering their adhesive and migratory behaviour. We show that these results are consistent with a haptotactic model of directed fin mesenchyme migration. Hence, Slit-Robo and S1P coordinate to provide tension to the interstitial matrix of the fin, thus driving robust tissue morphogenesis.

## RESULTS

### *stomp* mutant displays blisters in the caudal and pectoral fins

The *stomp* mutant was previously described as having variable degeneration of the pectoral fins (van Eeden et al., 1996). However, we noted this degeneration was preceded by formation of blisters in the pectoral fin fold (Figure 1A-D). We also observed small blisters in the caudal median fin in 40% of *sto* mutant embryos, suggesting *sto* affects all larval fins, as per other fin blister mutants (Figure 1E, F)(Carney et al., 2010). We noted that the penetrance of the *sto* phenotype was variable (Supplementary Table 1) as was expressivity, with 30% of *sto* mutants showing only unilateral pectoral fin blistering. H&E staining of coronal sections through the medial fin (Figure 1G, H) highlighted that blisters form in the proximal portion. The blisters form below the Laminin-positive basement membrane (Figure 1I, J), similar to that of Fraser-complex mutants (Carney *et al*., 2010). However, in contrast to the Fraser mutants, there was no loss of Fras1 protein at the basement membrane of blisters in *stomp* mutants (Figure 1K, L). The blisters that form in the fins of *sto* mutants are transient and collapse during later fin fold growth. We conclude that *stomp* represents a novel component required for fin integrity.

**Figure 1.**
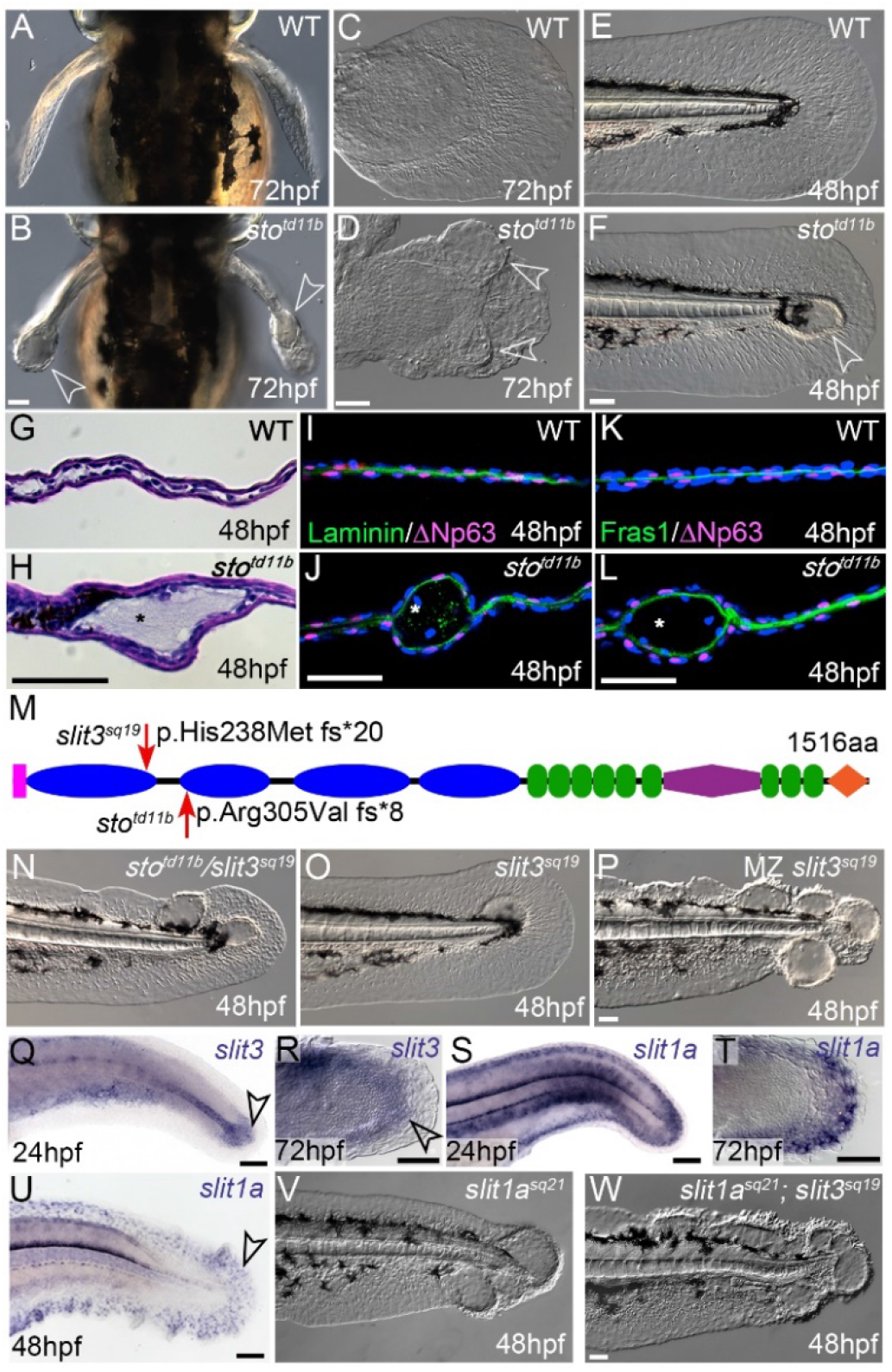
The *stomp* fin blister mutant corresponds to mutations in *slit3*. **A-F:** Dorsal (A,B) and lateral (C-F) images of the 3dpf pectoral (A-D) and 2dpf tail fins (E-F) of WT (A, C, E) and *sto^td11b^* homozygous mutants (B, D, F). Open arrowheads indicate blisters. **G-H:** H&E staining of coronal cryosections through the tail fin region of WT (G) and *sto* ^*td11b*^ mutant (H) embryos at 2dpf, **I-L:**Coronal confocal sections of tail fins from 2dpf WT (I, K) and *sto^td11b^* mutants (J, L), immunostained for ΔNp63 (I-L; magenta), Laminin (I, J; green) or Fras1 (K, L; green) and counterstained with DAPI (blue). Asterisks indicate blister cavity, which is below ΔNp63 positive basal keratinocytes and basement membrane labelled with Laminin and Fras1. **M:** Schematic of the zebrafish Slit3 protein, indicating the signal peptide (pink), four N-terminal domains with leucine-rich repeats (LRR, blue), six EGF-like domains (green), a lamininG domain (purple), three EGF-like repeats (green), and a C-terminal cysteine rich knot (orange). Location and nature of the *sto^td11b^* ENU and *slit3^sq19^* TALEN alleles are indicated at red arrows. **N-P:** Lateral Nomarski images of *slit3^td11b/sq19^* compound heterozygous (N), zygotic *slit3^sq19^* homozygous (O), and Maternal-zygotic (*MZ) slit3^sq19^* (P) tail fins at 48hpf. **Q-U:** Lateral brightfield images of tail (Q, S, U) and pectoral (R, T) fins stained by whole mount in situ hybridization for *slit3* (Q, R) and *slit1a* (S-U), indicating expression in proximal mesenchyme (arrowheads). **V-W:** Lateral Nomarski images of the *slit1a^sq21^* mutant (V) and *slit1a^sq21^*;*slit3^sq19^* double mutant (W) tail fins at 48hpf, indicating partial redundancy of Slit1a and Slit3 in tail fin morphogenesis. Scale bars: 50μm

### The *stomp* locus encodes *slit3*

We mapped *sto* to Linkage Group 14, refining to an interval containing 11 genes (Supplementary Figure 1A). Sequencing the coding region and intron-exon boundaries of 10 of these genes showed no plausible genetic lesion. However, a T to A transversion was found 7 bases upstream of the intron 9-exon 10 splice-site of the *slit3* gene (NM_131736; c.1341-7T>A; Supplementary Figure 1B, D). This was predicted to generate a novel splice acceptor, and sequencing *slit3* from *sto* mutant cDNA showed inclusion of 5 nucleotides from the end of intron 9 in the mature mRNA with the frame shift introducing 8 erroneous amino acids followed by a premature stop codon (c. 1340_1341insTGTAG; Supplementary Figure 1C, D). This truncates the 1516aa Slit3 protein at 305aa (Figure 1M). We noted that the new cryptic splice acceptor was not strong and sequence of *slit3* cDNA from homozygous *sto* mutants showed a mix of aberrant and correctly spliced transcripts. Therefore, to confirm that loss of Slit3 was responsible for fin blistering, we injected a translation blocking Morpholino against *slit3* embryos, which showed blistering in both the caudal fin and pectoral fins (Supplementary Figure 2A-C). Additionally, we used TALENs to create a frame-shifting indel mutation in exon 8 (Supplementary Figure 1E, F), which is predicted to lead a premature stop codon (*slit3^sq19^*; Figure 1M). This allele failed to complement *sto*, and 112 of 273 zygotic mutants of this allele showed tail blisters (Figure 1N, O).

RT-PCR showed *slit3* is expressed at all stages through to adulthood, including at the 2-cell stage indicating maternal contribution (Supplementary Fig 2D). We confirmed these observations by *in situ* hybridisation (Supplementary Fig 2E-F). We generated maternal zygotic *slit3* mutants and these had more severe tail fin blisters than zygotic *slit3^sq19^* mutants (Figure 1P). *In situ* hybridisation localised *slit3* expression to the proximal mesoderm region of both the tail and pectoral fins, from which the immigrating mesenchyme originate (Figure 1Q, R and Supplementary Fig 2G, H)(Lee et al., 2013). We also observed expression of *slit1a* in the larval tail and pectoral fins. *slit1a* expression remained in this population after invading the fin, whereas *slit3* was not expressed in the migrating mesenchyme (Figure 1S-U and Supplementary Fig 2I, J). Neither *slit1b* nor *slit2* were expressed in the posterior mesoderm of the tail which gives rise to the fin mesenchyme, although there was some expression of *slit2* in the proximal pectoral fin (Supplementary Figure 2K-N). We generated *slit1a* mutants through CRISPR/Cas9 mediated mutagenesis (*slit1a^sq21^*; Supplementary Fig 2O-R). Incrosses of *slit1a*heterozygotes gave 22.5% larvae with strong fin blisters at 48hpf (Figure 1V). Double *slit1a*; *slit3* zygotic mutants had more severe blisters compared to either mutant, indicating functional redundancy (Figure 1W), whilst *slit1a*^+/-^ crossed to *slit3* ^+/-^ gave clutches with 17.7% (n=388) of larvae having blisters.

### Robo receptors are required for Slit3-mediated fin morphogenesis

Slit proteins signal through Robo receptors and also bind a number of ECM components (Hu, 2001; Xiao et al., 2011). *In situ* hybridization revealed that Robo receptors are expressed in the fin fold, in a complementary pattern to that of the Slit ligands, with all three *robo* receptor genes (*robo1*, *robo2*, and *robo3*) dynamically expressed in the apical and sub-apical ectodermal ridge cells of the developing fin folds at different stages (Figure 2A-H; Supplementary Figure 3A-F). Subsequently, we investigated fin morphology in the zebrafish mutants for *robo2* (*ast^te284^*) (Fricke et al., 2001) and *robo3* (*twi^tx209^*) (Burgess et al., 2009). These mutants alone, or as double mutants, showed no fin defect (Figure 2J). We generated a TALEN-mediated knockout of *robo1* (*robo1^sq20^*;Figure 2I; Supplementary Figure 3I-K;) and although mild blistering was apparent in the pectoral fin of 13 of 28 (46%) *robo1* mutants at 72hpf (also seen with a *robo1* Morpholino (MO); Supplementary Fig 3G-H), there was no apparent tail fin blistering, either alone or combined with *twi* mutants (Figure 2K). As these genes are closely linked, to make triple deficient embryos we resorted to injection of *robo1* or *robo2* morpholinos into *ast;twi* or *robo1;twi* double mutants, respectively. Pronounced epidermal blistering was observed in both cases (Figure 2L, M). This indicates that Slit proteins function through their canonical receptors in maintaining integrity of the forming fin and that there is redundancy among Robo receptors in this function. As the only common expression domain of all three Robo receptors is the AER, we conclude that Slits within the developing fin fold are signalling to the AER cells.

**Figure 2.**
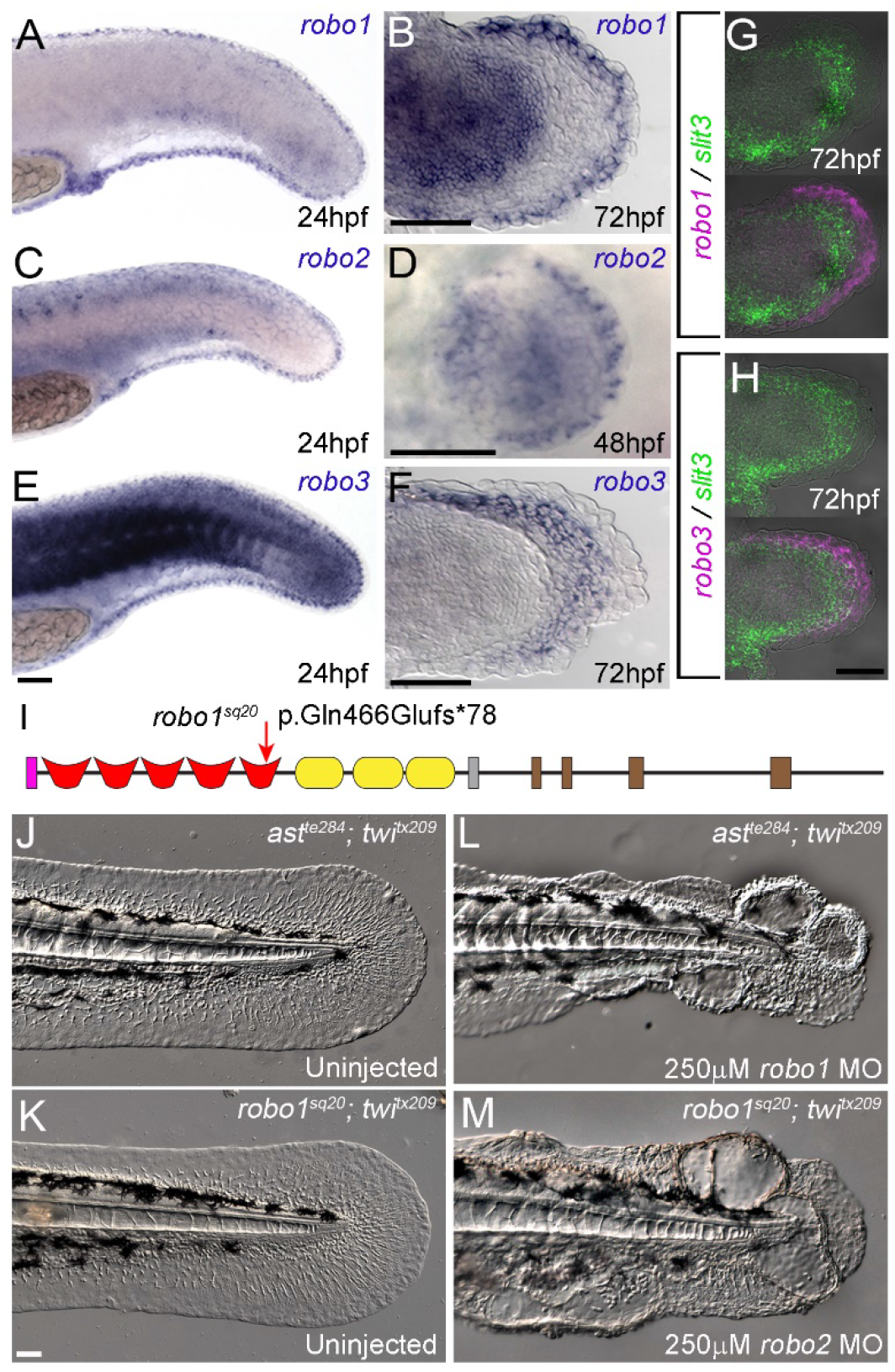
Robo receptors are expressed in the AER cells and act redundantly in fin morphogenesis. **A-F:** In situ hybridisation of tail (A, C, E) and pectoral (B, D, E) fins at 24, 48hpf or 72hpf, using probes for *robo1* (A-B), *robo2* (C-D), and *robo3* (E-F). Expression is seen in the apex of the fins, and/or sub-apically in the fins for *robo3*. **G-H:** Double fluorescent in situ hybridisation of 72hpf pectoral fins for *slit3* in green with either *robo1* (G) or *robo3* (H) in magenta. **I:**Schematic of the zebrafish Robo1 protein, with the position and nature of the TALEN-induced *robo1^sq20^* lesion. Domains shown are signal peptide (pink), five immunoglobulin (Ig) motifs (red), three fibronectin type III (Fn III) motifs (yellow), a transmembrane domain (grey), and cytoplasmic domains (CC0-3; brown). **J-M:**Lateral Nomarski images of tail fins of 48hpf larvae with double homozygous mutations in *robo3* (*twi^tx209^*) combined with either *robo2* (*ast^te284^*) (J, L) or *robo1* (*robo1^sq20^*) (K, M). Larvae were uninjected (J, K) or injected with 250μM morpholino targeting *robo1* (L), or *robo2* (M). Triple deficient larvae (L, M) show significant blistering of the fin fold compared to uninjected double mutant controls (J, K). Scale bars: 50μm

### Slit-Robo pathway synergises with S1P signalling

We hypothesised that Slit3 acts with other pathways known to cause fin blistering. We previously showed that Fras1 immunoreactivity is not disrupted in *sto* mutant fins (Fig. 1L). In addition, there was no obvious loss of expression of any genes previously associated with fin blisters (Supplementary Fig. 4). The cardia bifida mutant *miles apart* (*mil*) also displays fin blisters (Figure 3A) and corresponds to mutations in the gene encoding sphingosine-1-phosphate receptor 2 (*s1pr2*) (Kupperman *et al*., 2000). Although the hearts of *slit1a*; *slit3* mutants developed normally (data not shown), we noted similarity between the fin defects of *s1pr2* and *slit3* mutants (Fig 3A, B). To test for synergy between the two signalling pathways, we crossed *s1pr2* and *slit3* heterozygotes, to create *s1pr2*^+/*te273*^; *slit3*^+/*sq19*^ trans-heterozygotes. Depending on the clutch, between 2.5 to 25% of these showed genetic interaction, presenting with tail fin blisters (Figure 3E), never seen in the respective heterozygotes (Figure 3C, D). In addition, a low frequency of *slit1a*^+/*sq21*^; *s1pr2*^+/*te273*^ transheterozygotes also showed mild blistering of the fin Figure 3F; 5.1% (4/79) of trans-heterozygotes). Generation of trans-heterozygotes between *sto^td11b^* and four of the Fraser-class blistering mutants (*bla_ta90_*; *nel^tq207^*; *pif^te262^*; *rfl^tc280b^*) failed to display any genetic interaction, nor did *pif*^+/*te262*^; *s1pr2*^+/*te273*^ transheterozygotes (Supplementary Figure 5). Gα_13_ is an established downstream effector of S1pr2, and reduction of both Gα_13_ paralogues by Morpholino injection results in cardia bifida and tail fin blistering in zebrafish embryos (Ye and Lin, 2013). Injection of 200μM of the *gna13α* morpholino alone into WT embryos showed no fin morphology defect at 48hpf, however injection of the *gna13α* morpholino into *slit3*^*sq19*/+^ heterozygotes produced extensive fin blistering (Figure 3G-H). Thus, reduction of S1P pathway activity at two levels - by either genetic mutation or morpholino - demonstrates interaction with the Slit-Robo pathway.

**Figure 3.**
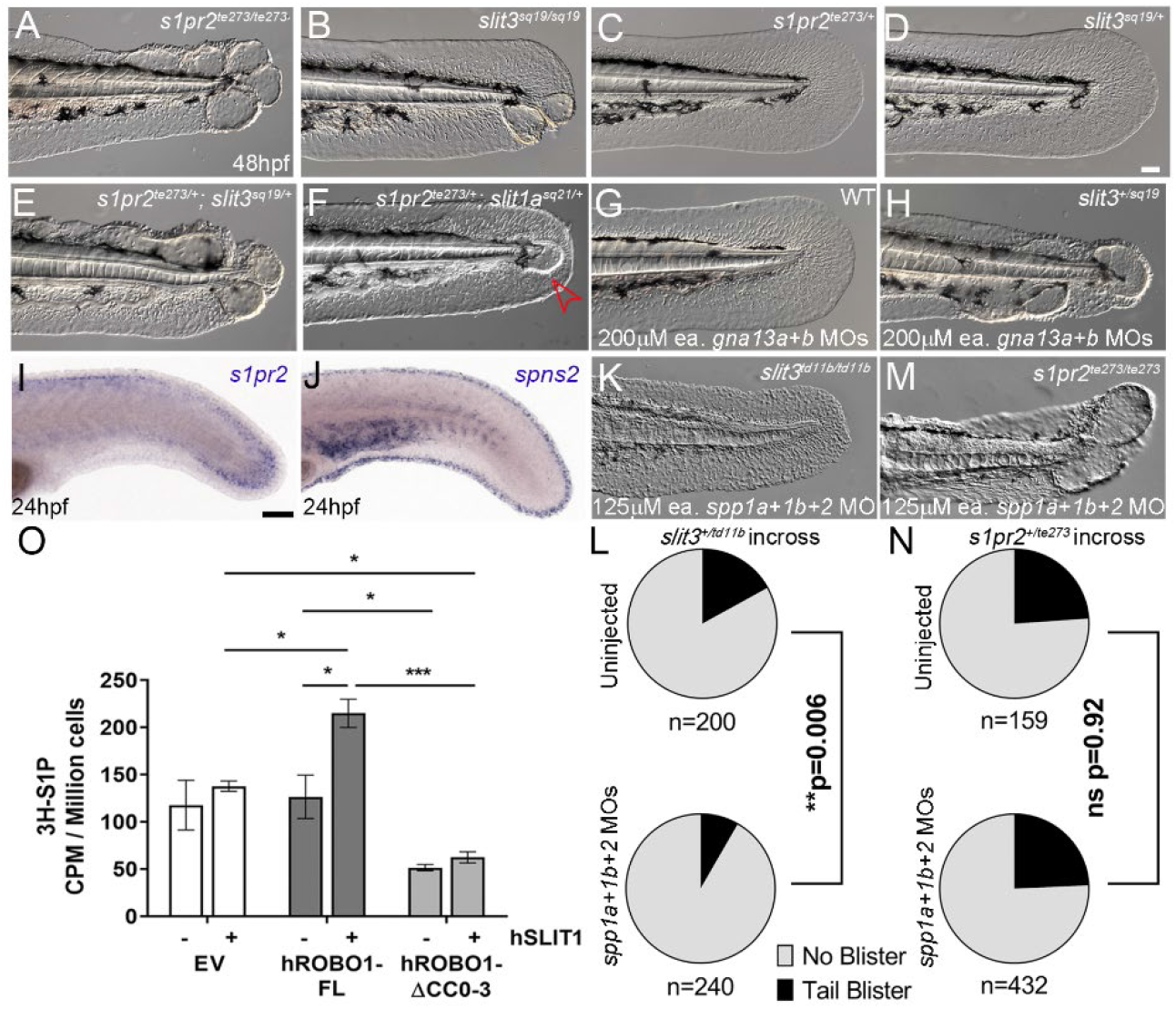
Sphingosine-1-phosphate signalling acts downstream of Slit-Robo signalling. **A-F:** Lateral Nomarski images of 48hpf tail fins of *s1pr2^te273/te273^* (A) and *slit3^sq19/sq19^* (B) homozygous mutants, *s1pr2*^+/*te273*^ (C) and *slit3*^+/*sq19*^ (D) heterozygotes, *s1pr2*^+/*te273*^; *slit3*^+/*sq19*^ (E) and *s1pr2*^+/*te273*^; *slit1a*^+/*sq21*^ compound heterozygotes. Blister in the compound heterozygotes highlighted with red arrowhead (E). *G-H:* Lateral Nomarski images of 48hpf tail fins of WT (G) and *slit3*^+/*sq19*^ heterozygous larvae injected with 200μM each of morpholinos against *gna13a* and *gna13b*.**I-J:** In situ hybridisation of tail fins at 24hpf using probes detecting the reciprocal expression of *s1pr2* in the emerging mesenchyme (I) and *spns2* in the apical cells (J). **K-L:** Morpholino reduction of the S1P catabolic enzymes *spp1a*, *spp1b*, and *spp2* rescues fin blistering of *slit3* mutants. Lateral Nomarski images of tail fins of *slit3^td11b^* (K) and *s1pr2^te273^* (M) mutants at 48hpf, which are injected with 125μM Morpholinos against each of each of *spp1a*, *spp1b* and *spp2*. **L,N:** Proportion of larvae derived from *slit3*^+*td11b*^ (L) or *s1pr2*^+/*te273*^ (N) heterozygous incrosses, with WT (grey) or blistered (black) fins, and injected with 125μM of Morpholinos against *spp1a*, *spp1b* and *spp2* (lower charts) or uninjected (upper charts). Significant reduction of larvae with blisters was seen between morpholino injected and uninjected clutches from *slit3*^+/*td11b*^ incrosses but not *s1pr2*^+/*te273*^ incrosses (Chi-squared test). **O:** HaCat cells over-expressing Robo1 (dark grey bars), truncated Robo1 (light grey bars), or vector control (white bars) were metabolically labelled with ^3^H-sphingosine. Cells were stimulated with recombinant SLIT1 (+) or unstimulated (-). Radiolabelled extracellular S1P was measured by scintillation counting and corrected for cell number. Means ± SEM shown; n = 3-4, * *p*<0.05, ** *p*<0.01, *** *p*<0.005, **** *p*<0.001 as determined by student t-test. Scale bars: 50μm

We additionally tested if *slit3* heterozygous larvae were sensitive to reduced S1pr2 signalling through use of the S1PR2 modulator, CYM-5478 (Satsu et al., 2013), which appears to inhibit S1pr2 in zebrafish and induces fin blisters in *s1pr2*^*te273*/+^ embryos in a dose dependant manner (Supplementary Figure 6A-C). 100% of embryos derived from a *slit3*^+/-^ x *slit3*^-/-^cross treated with 10-50μM CYM-5478 displayed fin blisters, as compared to the expected 45% untreated crosses (Supplementary Figure 6D-F). Similarly, treatment of embryos from a *slit3*^+/-^ outcross with CYM-5478 invoked fin blistering in a dose dependant manner (Supplementary Figure 6G). Genotyping indicated embryos with blisters were significantly more likely to be *slit3* heterozygotes (chi-squared; p<10^-4^; Supplementary Figure 6H). Thus CYM-5478 acts as an S1pr2 antagonist in zebrafish and synergises with *slit3* and *s1pr2* heterozygosity, providing further evidence of Slit-Robo-S1P signalling cross-talk in maintaining fin integrity.

*In situ* hybridisation revealed that *s1pr2* is expressed in the mesodermally derived fin mesenchyme, whilst the S1P transporter, *spns2*, is expressed in a complementary manner at the AER (Figure 3I-J). This indicates that the AER cells are the likely cellular source of S1P within the fin fold. Given that Robo receptors are also found in the S1P-producing cells, whilst S1pr2 is expressed in Slit-ligand-expressing mesenchyme, this suggests the interaction of the pathways is sequential and not due to parallel functions. This leads to the prediction that one pathway might regulate generation of the other pathway’s ligand.

### Slit-Robo pathway promotes S1P signalling

We tested if S1P production is epistatic to Robo function in two ways. We attempted to increase S1P levels in *slit3* mutants, by blocking S1P dephosphorylation. We injected three Morpholinos targeting the S1P phosphatases (*spp1a*, *spp1b*, *spp2*) into embryos derived from *slit3*^+/*td11b*^ incrosses. Whilst 17% (n=200) showed blistering in uninjected clutches, combined injection of *spp1a*, *spp1b* and *spp2* MOs resulted in a significantly lower frequency of blistering (8%, n = 240, p< 0.01; Figure 3K, L). Notably, when we genotyped all embryos with normal fins, the number of morphologically normal embryos with *slit3^td11b/td11b^* genotype was significantly higher (p < 0.005) in the *spp* MOs-injected group (22.5%; 18 of 80) compared to uninjected control group (5%; 4 of 80), suggesting partial rescue (Supplementary Figure 7A). In parallel, we injected the *spp* MOs into offspring of *s1pr2*^+/*te273*^ incrosses, but did not observe any rescue (Figure 3M, N) and found no increased representation of *s1pr2^te273^* homozygotes in phenotypically normal larvae injected with the *spp* Morpholinos (Supplementary Figure 7B). Taken together, reducing S1P dephosphorylation cannot compensate for loss of S1pr2, but can rescue loss of Slit3. We interpret this as indicating that Slit-Robo signalling lies upstream of S1P, through regulation of S1P generation or its release.

To investigate further if activation of the Slit-Robo pathway alters production and/or release of S1P, immortalised human keratinocytes HaCaT cells were transfected with tagged versions of full-length human ROBO1 (hROBO1-FL), a dominant negative truncated hROBO1 lacking the cytoplasmic domain (hROBO1-ΔCC0-3), or an empty vector, and expression confirmed by immunoblotting (Supplementary Figure 7D). After labelling of cells with ^3^H-Sph, levels of both intracellular and extracellular S1P were measured by scintillation counting. Expression of full-length ROBO1 receptor had no significant effect on extracellular S1P levels, whilst there was a slight increase in intracellular S1P upon ROBO1 expression (Figure 3O; Supplementary Figure 7C). We then stimulated these cells with recombinant hSLIT1, which resulted in a significant increase in intracellular and extracellular S1P in cells expressing the full-length hROBO1 (Figure 3O). Expression of the truncated hROBO1-ΔCC0-3 receptor significantly reduced extracellular S1P levels, compared to cells transfected with empty vector or ROBO1-FL in both stimulated and unstimulated conditions (Figure 3O). Curiously, intracellular levels of S1P were also significantly increased upon expression of hROBO1-ΔCC0-3 when rSLIT1 was supplied (Supplementary Figure 7C). These results suggest SLIT-ROBO signalling promotes synthesis and release of S1P in human keratinocytes.

### S1P establishes fin mesenchyme elongation and polarity

With the S1pr2 receptor expressed on mesenchymal cells, we hypothesised that a common defect in mesenchyme behaviour and function would account for the blistering in both *mil* and *sto* mutants. We crossed both mutants to the enhancer trap line, *sqet37Et*, which labels fin mesenchyme (Lee *et al*., 2013), to visualise tissue and cell morphology and behaviour. 3D visualisation indicated large blisters form in both mutants and that the mesenchymal cells remain attached to the inner wall of the epidermis (Figure 4A-C; Supplementary Figure 8A and Supplementary Movie 1). Timelapse imaging revealed that in the absence of either *slit* ligands or *s1pr2*, distinct blisters emerge around ~30hpf, continue to enlarge over the next several hours, until collapsing (Supplementary Movie 2, Supplementary Figure 8B). These movies indicated that mesenchymal cells migrate towards the AER, but stop just before reaching it, whilst more proximal cells form a tiled pattern behind. These cells normally have polarised morphology with a proximally positioned cell body and nucleus. They typically have one to three long directional protrusions orientated towards the distal fin tip (Figure 4A, G; Supplementary Figure 8B). Such protrusions are particularly prevalent for the tier of mesenchyme nearest the apex. The elongation increases over time as the cells migrate distally, such that in wild type embryos, mesenchymal cells reduce their circularity, increase eccentricity and elongate as they approach the periphery between 24hpf and 40hpf. In both mutants, mesenchymal cells maintain their circularity throughout, fail to increase either their eccentricity or their length (Figure 4D-I, Supplementary Movie 2, 3).

**Figure 4:**
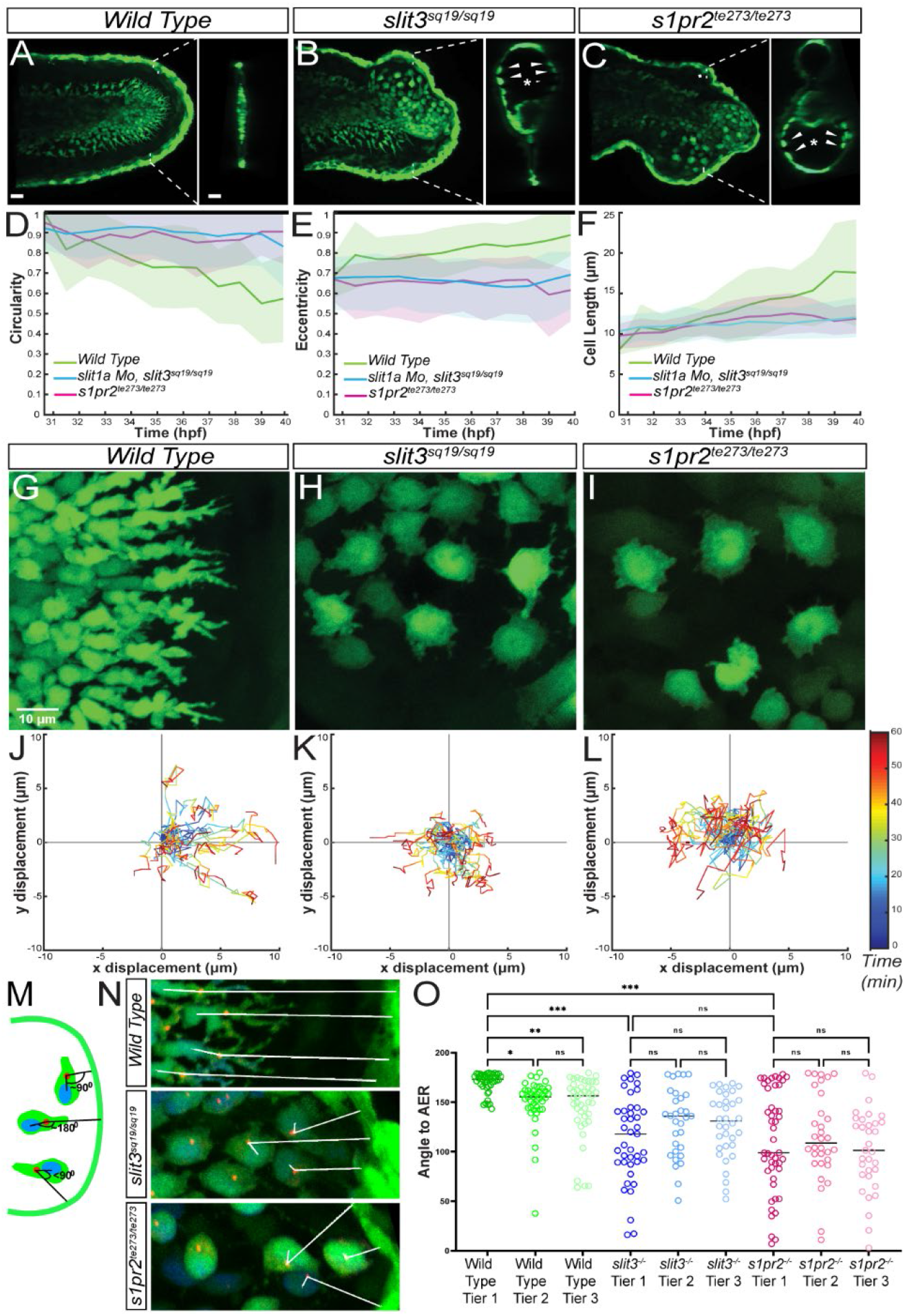
Mesenchymal cells of both mutants show abnormal morphology and loss of polarity. **A-C:** Confocal projections of the 40hpf tails of WT (A), *slit3^sq19^* (B) and *mil^te237^* (C), crossed to *sqet37Et*, labelling the fibroblasts in eGFP. Insets show transverse orthogonal slice at the indicated location. The mutant mesenchymal cells are attached to the inner wall of the blister as indicated by arrowheads. Scale bar indicates 20μm. **D-F:** Changes in fibroblast circularity (D), eccentricity, and length (E) as the cell migrates away from the paraxial mesoderm, between 30hpf-40hpf. Three embryos were tracked for each condition, and 33 cells of WT, 33 cells of *slit3*^*sq1*9^ and 35 cells of *mil^te237^* were analysed. **G-I:** WT cells (G) close to the apex have an elongated and polarised appearance whilst both mutants are unpolarised and have a disc like appearance (H,I, centre and right panels). **J-L:** Tracks of cells from WT (J) *slit3*^-/-^ (K) and *mil*^-/-^ (L) embryos over 60 minutes duration. Mutants display a lack of directionality and reduced displacement over a short range. Tracks are normalized to a common start point, 23 cells of WT, 22 cells of *slit3^sq19^* and 21 cells of *mil^te237^* from 3 to 4 embryos were tracked. **M-O:** Schematic describing the measurement of a cell’s approach angle to the nearest point on the AER (M). Merged, immunofluorescent images of WT (top), *slit3^sq19^* (centre) and *mil^te237^* (below) mesenchymal cells in *sqet37Et* background stained for EGFP (green), Gamma tubulin (red) and DAPI (blue). White lines run from the centre of nucleus to the nearest point on the AER, through the MTOC (N). Graph depicting the approach angles to AER of leading (Tier 1), following (Tier 2) and trailing (Tier 3) cells of WT, *slit3*^*sq1*9^ and *mil^te237^* embryos (O). A minimum of 30 cells were measured for each tier of each genotype. Scale Bars: 50μm (A), 10μm (G).

High resolution tracking of WT mesenchyme during migration indicated that these cells exhibited active filopodia directed towards the outer fin edge and directional movement away from their proximal origin (Figure 4G and J, Supplementary Movie 4). In contrast, mesenchyme positioned in the developing blisters of *slit3* or the *mil* mutants showed a discoidal morphology with multiple small, active yet short lived protrusions. These protrusions rapidly retracted and occurred in all directions around their periphery, implying impaired polarity (Figure 4H-I, Supplementary Movie 4). Indeed, migratory tracks of such cells in mutant embryos exhibited no directional preference and an overall reduced displacement towards the AER (Figure 4K-L, Supplementary Figure 8C-D).

We determined the orientation of the cells towards the AER, by measuring the angle from the nucleus through the MTOC (marked by γ-tubulin) to the nearest point of the AER (Figure 4M). In WT embryos, cells closest to the AER (Tier 1 cells) were the most polarised in the direction of migration, with angle to AER almost always close to 180°, with cells further from the AER less orientated towards the periphery (Figure 4N, O). In contrast, MTOC’s in all mesenchyme of both *slit3* and *mil* mutants were orientated far more randomly with respect to the nucleus and the nearest point on the AER (Figure 4N, O).

We conclude that whilst mesenchyme in both *slit3* and *mil* mutants adhere to the inner surface of the fin epithelium, they fail to polarise or generate productive filopodia, and do not correctly migrate towards the AER.

### S1P is required for stress fibres in mesenchyme

The fin malformations developed below the basement membrane and initiated around the mesenchyme. Thus, we hypothesised that the altered mesenchymal cell morphology and blistering in both mutants results from loss of cytoskeleton organisation or cellular adhesive mechanisms, such as focal adhesions and stress fibres. Indeed, S1PR2 signalling is well known to induce stress fibre formation and inhibit cell migration via Gα_12/13_ activation of PDZ-RhoGEFs (Yamamura et al., 2000). Further, either suppression of Gα_13_ expression or injection of a dominant negative form of Arfgef11 (a PDZ-RhoGEF) results in tail blisters (Ye and Lin, 2013). To visualise cellular focal adhesions and stress fibres, we performed immunofluorescent staining for phospho-focal adhesion kinase (pFAK) and phospho-non-muscle myosin II (pNM-myosin II), respectively. WT fin mesenchyme had strong staining of both pFAK and pNM-myosinII localised to the distal protrusions of the most apical cells (Figure 5A,A’,B,B’). Strikingly, there was a gradient of signal across the fin with high signal apically and little to no signal in proximal mesenchyme. Both *mil* and *slit3* mutant mesenchyme lacked any evidence for pFAK or pNM-myosin II, irrespective of location in the fin fold (Figure 5A,A’,B,B’).

**Figure 5:**
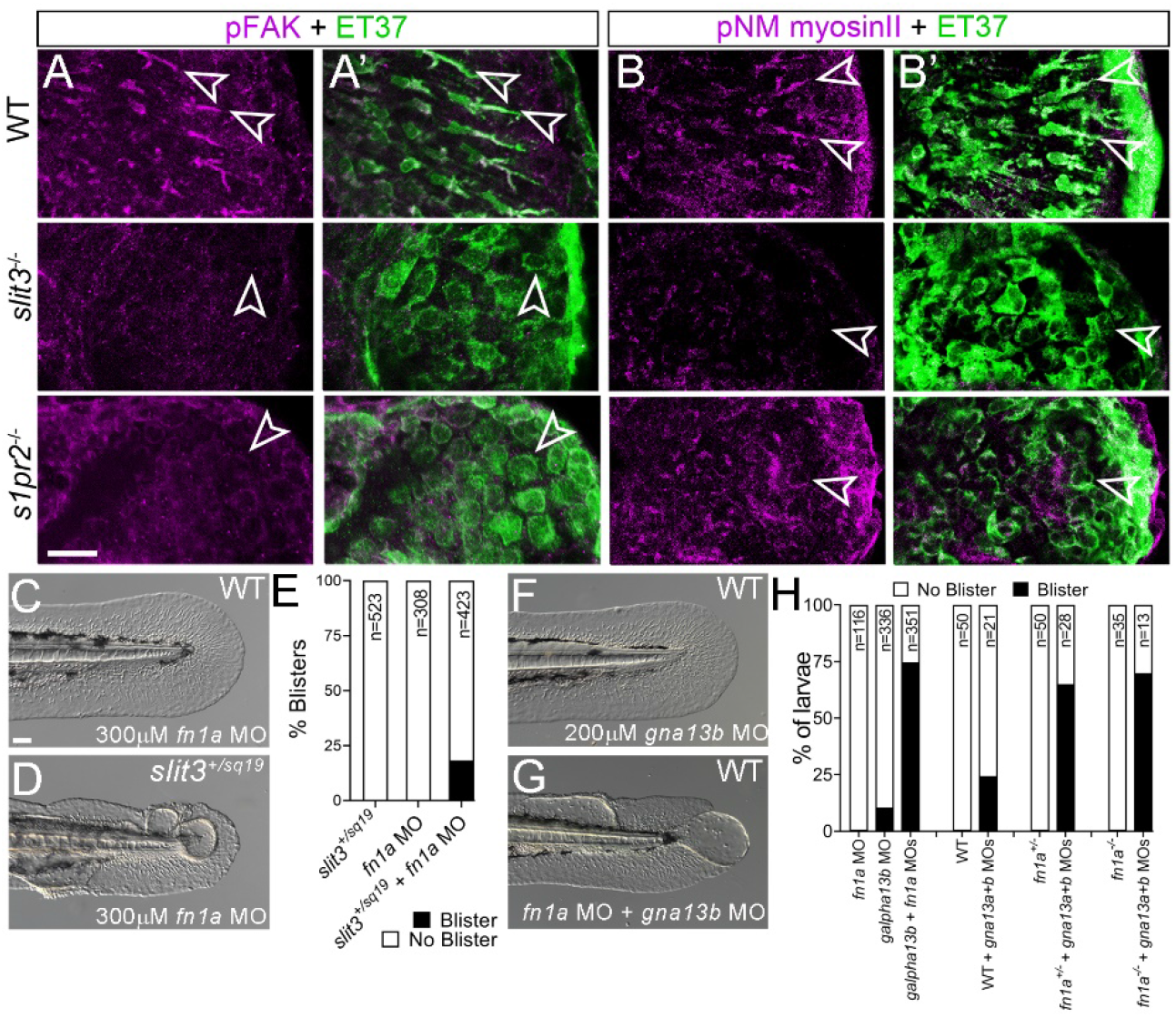
Both *slit3* and *mil* mutants show loss of focal adhesion markers and sensitivity to Fibronectin levels. **A-B:** Immunofluorescent staining of 48hpf tail fins in WT *sqet37Et* (Top row), *slit3^sq19/sq19^*; *sqet37Et* (middle row), and *s1pr2^te273/te273^*; *sqet37Et* (bottom row) transgenic larvae, stained for phospho-FAK (magenta; A, A’), phospho-Non Muscle myosin (magenta; B, B’) and eGFP (green; A’, B’). Mutant mesenchymal cells show significantly reduced p-FAK and p-NM myosin II (magenta; A, B) signals in *slit3* and *s1pr2* homozygous mutants compared to the WT fin mesenchyme. Arrowheads indicate fin mesenchymal cells in the larval fins. **C-E:**Lateral Nomarski images of 48hpf larval fins which are WT (C) or *slit3*^+/*sq19*^ heterozygotes (D) and are injected with 300μM *fn1a* morpholino. Blisters are observed only when there is reduced Fn1a in *slit3* heterozygotes (quantified in **E**). **F-G:** Lateral Nomarski images of 48hpf WT larval fins injected with 200μM *gna13b* alone (F) or with 300μM *fn1a* morpholino (G). **H:** Quantification of the proportion of larvae with fin blisters when low amounts of *gna13* morpholinos (125μM each *gna13a* and *gna13b* MO combined or 200μm *gna13b* MO alone) are injected into WT, *fn1a* morphants (300μM MO), *fn1a*^+/-^ heterozygotes and *fn1a*^-/-^ mutants. Loss of a single or both copies of *fn1a* exacerbates reduced *gna13* levels, as does knockdown of *fn1a*. Scale bar A-B: 20μm; C-G: 50μm.

Fibronectin1a (Fn1a) (Trinh and Stainier, 2004), in concert with Spns2 (Hisano et al., 2013), S1pr2 (Matsui et al., 2007), and Gα_13_ (Ye et al., 2015), is required for the migration of myocardial precursors. Whilst the fins of most *fn1a* mutants appear normal, the interaction of *fn1a* with *gα13* in cardiac migration suggests that they may interact during fin morphogenesis. Low doses of *fn1a* or *gna13b* MOs alone yielded no or rare fin blisters respectively, but following combined injection, 74% of larvae had distal fin blisters reminiscent of those in *sto* and *mil* (Figure 5C, F-H). Similarly, injection of non-phenotypic doses of *gna13a* and *gna13b* MO into *fn1a* mutants or heterozygotes significantly increased the proportion of larvae with blisters, compared to WT injected with these MOs (Figure 5G-H). Given that we have linked Slit-Robo signalling with the S1P – *gα13* pathway, we would expect that *sto* might interact with partial loss of Fn1a. Indeed, injection of low doses of *fn1a* MO into *slit3* heterozygotes realised about 18% of larvae with fin blisters (Figure 5C-E). Immunostaining for Fibronectin indicates it is localised to the fin fold interstitium, and that fibronectin protein remains in the fin dermis of both *slit3* and *s1pr2* mutants (Supplementary Figure S9A-C).

We thus propose that S1P is acting through S1pr2 and Gα_13_ to establish mesenchyme adhesion to Fn in the interstitial ECM. These mesenchymal cells indeed specifically express integrin receptors for fibronectin, Itgb3b and Itgav, which are known to promote fibroblast contractility on Fn substrates (Fiore et al., 2018) (Supplementary Figure 9D, E). However, attempts to ablate these proteins by morpholino yielded moderate gastrulation and axis defects and CRISPR mutants had no phenotype, suggesting compensation.

### Directed mesenchyme migration by a self-generated signalling gradient interacting with the fin boundary

Combining our above observations, we hypothesise that the S1P activated adhesion of the mesenchymal cells impart tension on Fibronectin in the interstitial ECM, retaining the two epidermal sheets of the fin fold in close proximity, whilst also promoting mesenchyme polarity and migration (Figure 6A).

**Figure 6.**
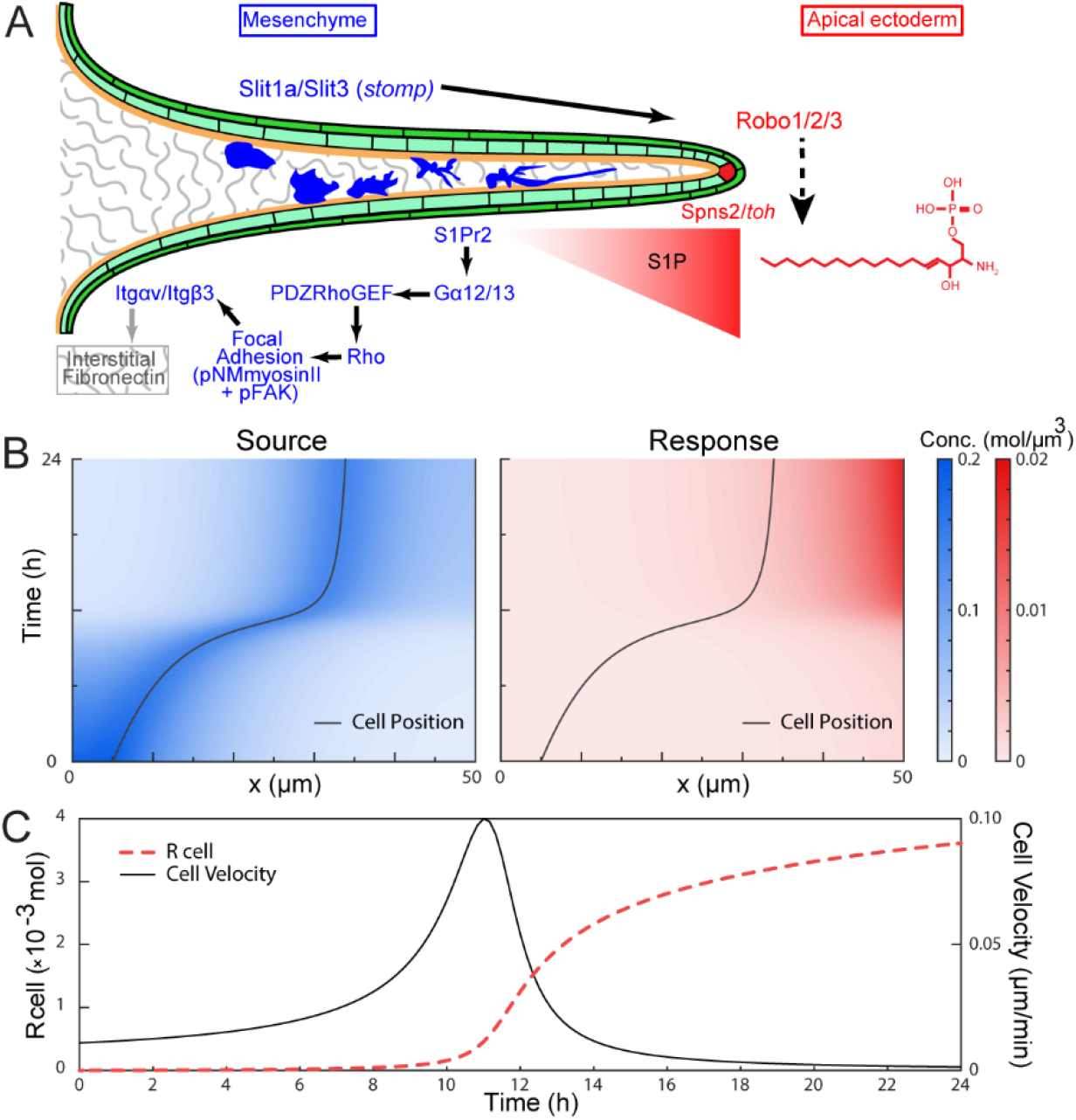
Reciprocal signalling of Slit-Robo and S1P creates an adhesion gradient that modulates cell migration. **A:** Model of Slit-Robo and S1P signalling deployment in the fin fold (light and dark green), with apical ridge cell in red and mesenchymal cells in blue invading the fin. Fibronectin of the interstitial ECM is in grey stipples. Components found in or generated by the mesenchymal cells are listed in blue, whilst those of the apical ridge cells are in red. The gradient of S1P (red) is shown as a triangle and the pathway activated by S1 PR2 in mesenchymal cells is shown in blue, culminating in adhesion to interstitial fibronectin (grey). Robo signalling promotes production and or release of S1P (dashed arrow). **B-C:** Computer simulation of a single mesenchymal cell migrating towards the apical ridge (x = 50μm) under reciprocal signalling. The cell emits a source signal, S, which induces the production of a response signal, R, from the apical ridge (B). The resulting cell velocity depends on the amount of R present at the cell position, R cell, with moderate levels of R cell resulting in the highest cell migration rates (C)

Why this reciprocal signalling between mesenchyme and apical ridge cells has been established in the fin is not clear. As S1P is released from a discrete source at the apex, it likely forms a gradient along the distal-proximal axis of the fin fold, and hence a gradient of adhesiveness, as seen by pFAK and pNM-myosin II staining. Proximity of Slit expressing mesenchyme to the apical domain might alter the level of S1P release, which would act to sharpen the adhesion gradient of the mesenchyme as it approaches the fin fold apex.

To test whether this idea is plausible to direct cell migration, we constructed a simple model of the interactions between the migrating cells, which secrete Slit, with a return gradient of S1P (Methods). Our simulation results suggest that such a mechanism may allow the cell to direct its own migration by interacting with the boundary to adjust its velocity as they migrate (Figure 6B-C). The cell is brought to a final position within the tissue environment when the adhesion strength prohibits further migrations. This mechanism enables fine tuning of the adhesion experienced by the cell as it migrates through the fin fold; in essence, cells can regulate the haptotactic field they encounter during migration to the apex and alter tissue shape as cells approach their destination.

## DISCUSSION

It has been established that there is a distal-proximal gradient of cell-cell adhesion in the forming limb bud, critical for correct morphogenesis (Wada, 2011). Whether cell-matrix adhesion also shows a gradient is not known. Additionally, limb bud mesenchyme polarity and migration are defined by AER derived signals such as Wnt5a, and that cell proximal-distal elongation drives limb morphogenesis (Gros *et al*., 2010; Wyngaarden *et al*., 2010). It has been proposed that the distal-proximal gradient of adhesion cooperates with orientated cellular behaviour for morphogenesis (Wada, 2011). Our work uncovers an unexpected role for the Slit-Robo pathway in the morphogenesis of the medial and paired fins of zebrafish, considered to be the evolutionary precursors of tetrapod limbs. In *slit3* mutants, fin mesenchyme has defects in polarity, stress fibre formation, Fibronectin adhesion, and migration leading to disrupted fin morphology. The tissue, cellular and molecular defects of *slit3* mutants are replicated in the fins of the *s1pr2* mutant, and we see synergy between the Slit-Robo and S1P signalling pathways by combined genetic and/or pharmacological disruption. Localisation of the receptors of the two pathways, as well as genetic epistasis analysis, supported a model of Robo signalling promoting generation or release of S1P from the fin AER. This was corroborated by *in vitro* S1P biochemical assays which also suggested this regulation occurs in mammalian cells. In turn, S1P is received by the immigrating mesenchymal cells, where the relevant receptor, S1pr2, is expressed. Activation of S1PR2 is described to induce stress fibres and focal adhesions via Rho (Wang *et al*., 1997), and we observe loss of markers of both these adhesive structures in both *slit3* and *s1pr2* mutants. Furthermore, we have seen that partial loss of components of the Slit-Robo or S1P pathways render larval fins sensitive to reduced levels of fibronectin. We hypothesise that the mesenchymal cells bind to interstitial fibronectin via their activated focal adhesion complexes and S1P activation of myosin in the stress fibres both promotes initial directional migration and also provides tension on the interstitial matrix of the most distal fin fold. It is plausible to consider that this tension retains the two epidermal sheets of the fin fold in close proximity. These results are summarised in Figure 6A.

Missense mutations in *S1PR2* have been found in three families with autosomal recessive hearing impairment (Hofrichter et al., 2018; Santos-Cortez et al., 2016). Intriguingly, for one of these families, all individuals with hearing impairment also had distal limb anomalies. As they were not seen in the other families nor the *S1pr2* mouse mutants, a role for S1PR2 in limb development was excluded, however no other mutations were identified that may account for these limb malformations, and the cause in this family remains unidentified. Given our identification of defects in mesenchyme morphology in *s1pr2* mutant fins, it may be worth revisiting a partially redundant role for S1PR2 signalling in human limb development.

How Robo signalling promotes secretion of S1P is unclear. We found *spns2* mRNA expressed at normal levels in *slit3* mutant fins and *slit3* is unlikely to act via *sphk2* transcriptional regulation as maternal *sphk2* alone is sufficient for normal fin formation (Mendelson *et al*., 2015). Robo receptors do not have enzymatic activity and, following binding by Slits, recruit activators to their intracellular domains. These include a number of actin cytoskeleton regulators including Slit-Robo GAPs (SrGAPs), Sos, and Pak (Blockus and Chedotal, 2016). We see co-expression of *srgap1a* and *srgap2* with the *robo* genes in the apical fin fold. However, combined morpholino knockdown of these *srgap* genes did not elicit a blister phenotype. It has been shown that Slit induces recruitment of Sos to the Robo receptor through promoting endocytosis of the ligand-receptor complex, and that Sos can access Robo only present in endosomes (Chance and Bashaw, 2015). In parallel, Shen et al have demonstrated that SPHK1 and SPHK2 both bind strongly to endocytic structures (Shen et al., 2014). However, our cell culture experiments, using overexpression of ROBO1 receptor and recombinant SLIT1, failed to show clear alteration of the sub-cellular localisation of SPHK2 or SPNS2.

Despite being mostly known for its role in axon guidance and neuron cell migration in both vertebrates and invertebrates (Jen et al., 2004; Kidd et al., 1999), a role for Slit-Robo signalling in morphogenesis is not novel. A patient with a translocation mutation affecting *ROBO2* has been described to have clinodactyly and syndactyly in addition to kidney and urinary tract defects (Lu et al., 2007), while a dominant *de novo* missense mutation in *SLIT2* was found in a patient with myopia and dermal connective tissue defects (Liu et al., 2018). Perturbation of Slit-Robo signalling leads to cardiac malformation in human, mouse, zebrafish and *Drosophila* (Fish et al., 2011; Kruszka et al., 2017; MacMullin and Jacobs, 2006; Mommersteeg et al., 2015). In the latter two species, Slit-Robo signalling is essential for migration of cardiac precursors to the midline (Fish *et al*., 2011; Santiago-Martinez et al., 2008). In particular, medially migrating endocardial cells in zebrafish *slit2* morphants show dynamic filopodia but lack directionality, reminiscent of the mesenchyme of the fins in *slit3* and *mil* mutants. Thus, both S1P and Slit-Robo signalling have been associated with cardiac precursor migration defects. Whilst we link the two pathways in fin morphogenesis, curiously the *slit3, slit1a* or *robo1* mutants did not show an overt defect in heart morphogenesis, despite all three showing distinct similarities with fin blisters in *miles apart*. It is possible that sub-functionalisation of *slit* genes has led to *slit2* functioning in the cardiac field whilst *slit1a* and *slit3* are important for fin morphology.

Examples of interaction of the Slit-Robo pathway with other cell signalling systems are limited (Blockus and Chedotal, 2016). Our work identifies a novel relay signalling system between the AER and the immigrating mesenchyme which is essential for cell-ECM adhesion, polarity and fin morphogenesis. This will refine biophysical models of how limb and fin outgrowth are constrained into precise morphologies.

## EXPERIMENTAL PROCEDURES

### Zebrafish strains and husbandry

Zebrafish were maintained in IMCB fish facility under standard conditions at 28°C on a 14 h light 10 h dark cycle. Embryos were obtained through natural matings, raised at 28°C in E3 medium (5mM NaCl, 0.17mM KCl, 0.33mM CaCl_2_, 0.33mM MgSO_4_), and staged according to Kimmel et al. (1995). The following lines were used: AB wild-type, *sto^td11b^*, *bla^ta90^*, *nel^tq207^*, *pif^te262^*, *rfl^tc280b^* (all described previously in van Eeden *et al*. (1996)), *mil^te273^* (Kupperman *et al*., 2000), *ast^te284^* (Fricke *et al*., 2001), *twi^tx209^* (Burgess *et al*., 2009) and the sqet37Et (ET37) enhancer trap line (Lee *et al*., 2013) in *slit3^sq19^* and *mil^te273^* backgrounds., *slit3^sq19^*, *robo1^sq20^* and *slit1a^sq21^* mutants were generated as described below. The *slit3^sq19^* mutation is a frame shifting indel, c.1141_1147delinsATG; p.His238MetfsTer20. The *slit1a^sq21^* mutation is a frame shifting indel, c.269_274delinsCCGACGCGCCGCGC; p.Ile90ThrfsTer15. The *robo1^sq20^* mutation is a 13bp deletion leading to a frame shift c.1396_1408del; p.Gln466GlufsTer78. All experiments were conducted under A*STAR BRC IACUC oversight (IACUC number 140924).

### Genetic mapping

For genetic mapping, *sto^td11b^* was crossed onto the WIK background and mutant and sibling offspring were each pooled for bulk-segregant analysis following Geisler (2002). This led to an assignment to linkage group 14. Fine single sequence linkage polymorphism mapping was then conducted on 430 single mutant embryos, placing the *sto* locus between z6847 and z22128. SNP markers were developed to refine the interval to a 1.1Mb interval. The coding regions and intron-exon boundaries for the 11 genes in that interval were sequenced and a mutation in *sto* larvae was identified in Intron 9 of the *slit3* gene.

### TALEN and CRISPR mutagenesis

Mutagenesis of *slit3* or *robo1* was performed by design, assembly and injection of TALEN constructs, which were made to target sites in exon 8 of each gene. For the *slit3* gene, the dimeric TALENs bound the following sites (5’-3’) in exon 8, Left: CACACAGTGCATGGCC; Right: CAGGGACATTGAGACC. For the *robo1* gene, the TALENs bound the following sites (5’3’) in exon 8, Left: CCACACATGATTCCCG; Right: CTGCAGGGCTCCAGTG. Repeat Variable Di-Residue (RVD) recognition modules for the above target binding sites were fused to the left or right monomer of the heterodimeric variant of FokI nuclease using the Golden Gate system as per Dahlem et al. (2012). Mutagenesis of *slit1a* was performed using the CRISPR-Cas9 system with the guide RNA targeting the exon 3 sequence 5’-GGAGAACCAGATTGTAACGG-3’. A PCR product containing a T7 promoter directly upstream of the sgRNA was generated using overlapping primers as per Bassett et al. (2013). TALEN and Cas9 RNAs were generated from plasmids linearised with NotI and synthesised with the mMessage Machine SP6 kit (Invitrogen) according to instructions. The *slit1a* sgRNA was synthesised from purified PCR product using the <u>MEGAshortscript^™^ T7 Kit</u> from Invitrogen as per manufacturer instructions.

Following injection of TALEN RNAs or*slit1a* CRISPR sgRNA with Cas9 RNA into wild-type embryos, a selection of larvae was sequenced to confirm efficient mutagenesis. The remaining larvae were raised to adulthood, incrossed and selected larvae sequenced for identifying founder adults carrying mutations.

### Morpholinos and Inhibitors

Morpholinos (MOs) used and their sequences (5’-3’) were as follows:

*slit3* ATG: CCCCCAATACTTTACCCACCGCATC; *robo1* ATG:
ATCCAATTATTCTCCCCGTCATCGT; *robo2* ATG:
GTAAAAGGTGTGTTAAAGGACCCAT; *spp1a* ATG:
ACCCCGCTTTTATCCCGCCTGCCAT; *spp1b* ATG:
ATCTGTGGAGCACGTCGCTTGCCAT; *spp2* ATG:
TCAGGTACGTGATGATTCTCCACAT; *fn1a* ATG:
TTTTTTCACAGGTGCGATTGAACAC; *gna13a* ATG:
AAATCCGCCATCTTTGTAGTAGCGA; *gna13b* ATG:
AGGAAATACGCCATCTTTGTGCAAC.

All MOs were obtained from GeneTools and dissolved to a stock concentration of 1mM in distilled water. For injection, stock MOs were diluted in 1X 1x Danieau’s solution: 5 mM HEPES (pH 7.6), 58 mM NaCl, 700 μM KCl, 400 μM MgSO_4_.7H2O, 600 μM Ca(NO_3_)_2_ with 0.5% phenol red and injected (125-500μM) individually or in combination into one-cell stage embryos.

S1pr2 selective modulatory agent, CYM-5478 (Aobious), was dissolved in DMSO as 25mM stock solution and added to embryos from 3hpf to 48hpf at final concentration of 10-50μM, and then scored for fin fold abnormalities.

### Microscopy and sectioning

Brightfield and Nomarski images were taken on a Zeiss AxioImager M2, whilst fluorescent images were taken on a Zeiss LSM700 confocal. A Zeiss LSM800 confocal was used for all timelapse confocal movies. Live embryos were mounted in 3% Methyl Cellulose for Nomarski images of the tail. For timelapse movies, embryos were anaesthetised in 0.02% tricaine buffered to pH7.0 and mounted in 0.7% Low Melting point agarose in glass bottomed imaging dishes. Embryos were then overlaid with 0.5xE2 medium (7.5mM NaCl, 0.25mM KCl, 0.5mM MgSO_4_, 75μM KH_2_PO_4_, 25μM Na_2_HPO_4_, 0.5mM CaCl_2_, 0.35mM NaHCO_3_) containing 0.02% tricaine (buffered to pH 7.0), and the agarose around the tail was excavated to permit free movement during growth.

For coronal sections, cryosectioning of embryos was performed using a Leica CM1900 cryostat and the 16μm sections were then stained by Haematoxylin & Eosin.

### Image Processing, cell shape analysis and tracking

All microscopy images were processed using Zen 3.1 software (Zeiss), Fiji (ImageJ, ver. 1.52p) or Imaris (Bitplane).

Images of developing zebrafish fins were aligned in 3D using a custom MATLAB code, and image segmentation was done using the surfaces function in Imaris 9.2.1. Quantification of the segmented data was done using the functions regionprops and regionprops3 in MATLAB.

Circularity, eccentricity and length of the cells, as they migrate away from the paraxial mesoderm, was measured on time-lapses (20x magnification), obtained between 30hpf-40hpf. The shortest Euclidean distance between the cell centroid and the paraxial mesoderm are measured and binned at 10um intervals. Within each distance interval, the mean and standard deviation of the circularity, eccentricity and length measures were calculated for cells of each condition. Three embryos were tracked for each condition, 33 cells for WT, 33 cells for *slit3^sq19^* and 35 cells for *mil^te237^*.

Cell circularity specifies the roundness of the object and is defined as 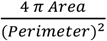 such that a perfect circle has a circularity value of 1. Cell eccentricity gives the elongation of the object and is defined as 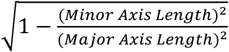 so that an ellipse with an eccentricity of 0 is a circle. The length of each cell is given by its major axis length.

Cell tracking was performed on time-lapse images (40x magnification), obtained between 36hpf-43hpf. The images were drift corrected with Imaris (Bitplane) to negate movement due to tissue growth, and further manually tracked using Fiji. The XY coordinates obtained were plotted using MATLAB.

A cell’s approach angle to AER was measured using the angle tool function of Fiji/ImageJ, with nucleus and MTOC as anchor points. Mesenchymal cells closest to the AER (most distally positioned) are considered Tier 1 cells. Cells positioned immediately behind Tier 1 cells are designated as Tier 2 cells. Tier 3 cells are positioned behind (proximal) the Tier 2.

### PCR, in situ hybridisation and antibody staining

Sequences for generating probes were amplified from cDNA by PCR using GoTaq DNA Polymerase (Promega) on a BioRad T100 Thermal cycler. Amplicons were purified using a Qiagen PCR purification kit, and then cloned into pGEMT-Easy (Promega). For the following probes, plasmids were linearised with *SacII* (NEB): *slit2*, *robo1*, *robo2*, *robo3*, *s1pr2*, *spns2*. The *slit1a and slit1b* plasmids were linearised with *ApaI*, whilst *slit3* probe plasmid was linearised with *MfeI*.For all RNA in situ probes, the SP6 DIG labelling kit (Roche) was used for transcription, except for *slit3* probe, which used either the T7 DIG or T7 Fluorescein labelling kits (Roche). Wholemount in situ hybridisation on embryos was performed as per Thisse and Thisse (2008), and developed using NBT/BCIP (Roche) and cleared in glycerol for imaging. Double fluorescent in situ hybridisation was performed using Fluorescein labelled *slit3* probe and DIG labelled *robo1* or *robo3* probes according to Brend and Holley (2009).

For immunofluorescent antibody stainings, embryos were fixed with 4% PFA for 2 hours room temperature, except for anti-pMLC2 and anti-pFAK stainings, which used 95% MeOH with 5% glacial acetic acid at -20°C for 4hrs. Embryos were permeabilised in Acetone for 7 mins at -20°C, washed in PBS with 0.5% Triton, blocked for 2 hours in Block solution (PBS Triton with 0.5% goat serum and 0.1% dimethyl sulfoxide), and then incubated in Block with primary antibody. After extensive washing in PBS Triton, embryos were incubated with secondary antibodies overnight in Block solution, and then rewashed in PBS Triton before clearing in glycerol for imaging. Primary antibodies, sources and dilutions used were as follows: mouse anti-ΔNp63 (Clone 4A4; Biocare, Cat# CM163; 1:500), rabbit anti-laminin (Sigma, #L9393, 1:200), rabbit anti-zebrafish Fras1 ((Carney *et al*., 2010), 1:50), rabbit anti-eGFP (Torrey Pines Biolabs, #TP401, 1:1000), ), rabbit anti-Fibronectin (1:200; F3648, Sigma-Aldrich), rabbit anti-phospho-FAK pY861 (1:250; #44-626G; Thermo Fisher Scientific), rabbit anti-phospho-Myosin Light Chain II (S19; pNM-myosin II) (1:250; #3671; Cell Signalling Technology) and rabbit polyclonal anti-Gamma tubulin (1:250, GTX113286, GeneTex). Secondary antibodies were sourced from Invitrogen and used at 1:400: Alexa 488-conjugated donkey anti-rabbit IgG, Alexa 546-conjugated donkey anti-mouse IgG, and Alexa 647-conjugated donkey anti-rabbit IgG. Counterstaining of nucleic was performed using 1μg/mL DAPI (Thermo Fisher Scientific).

### Generation of Robo1 expression vectors

Human Robo1 full length (FL) cDNA (GenBank accession number: NM_133631.3) was cloned with a C-terminal 3xHA tag into pcDNA3 from a hRobo1 ORF clone by PCR to generate hRobo1-FL-3xHA(C)/pcDNA3. The dominant negative truncated hRobo1 construct, hRobo1^ΔCC0-3^-3xHA(C)/pcDNA3, which included the first 920 amino acids (excluding the CC0-3 cytoplasmic domains) was PCR amplified from hRobo1-FL-3xHA(C)/pcDNA3 plasmid.

### Cell culture and S1P production assay

HaCaT cells were cultured in Dulbecco’s modified Eagle’s medium (DMEM) containing 10% fetal bovine serum and 100 units/mL penicillin and 100 μg/mL streptomycin in a 5% CO_2_ humidified incubator. The rate of S1P formation in intact cells was determined as an *in situ* assay of SphK activity as described previously (Zhu et al., 2017). Briefly, HaCaT cells were transfected with pcDNA3, hRobo1-FL-3xHA(C)/pcDNA3 or hRobo1^ΔCC0-3^-3xHA(C)/pcDNA3 using Lipofectamine2000 (Thermo Fisher Scientific) and incubated for 24 hours and then sub-cultured into 12-well culture dishes and allowed to bed down overnight. The cells were then labelled with 0.25 μCi of [^3^H]-sphingosine (Perkin-Elmer) in serum-free DMEM with 0.1% fatty-acid free BSA with and without the addition of 10ug/ml recombinant Slit1 protein. After 30 min incubation at 37°C in a humidified atmosphere of 5% CO_2_, the conditioned medium was removed and the cells washed and scraped into cold PBS. [^3^H]-S1P formed during the 30 min incubation was then extracted from both the conditioned medium and cell pellets via a modified Bligh-Dyer extraction. Briefly, 300 μl of acidified methanol (100:1, methanol: concentrated HCl) was added to the cell pellets and then sonicated for 30 s in an ice-bath. To each cell sample 300 μl of 2M KCl, 300 μl of chloroform, and 30 μl of 3M NaOH were then added. After vigorous mixing and centrifugation at 13, 000 x g (5 min) a phase separation enabled separation of S1P in the upper aqueous methanol phase from sphingosine in the lower chloroform phase. The [^3^H]-S1P in the upper aqueous methanol phase was then analysed by scintillation counting (Microbeta, Perkin Elmer). Extracellular [^3^H]-S1P in the conditioned medium (500 μl) was analysed in the same manner with the addition of 500 μl of methanol, 500 μl of chloroform, and 50 μl of 3M NaOH. All analyses were performed in triplicates and corrected for total cell number.

### Mathematical Model

We model the reciprocal signalling for a single cell, with position *x_cell_*, migrating on a static one-dimensional spatial domain bounded by the notochord (*x* = 0) and the apical ridge (*x* = *L*). Let *S*(*x*, *t*) denote the concentration of a ‘Signal’ molecule secreted by the migrating cell, corresponding to the Slit. Let *R*(*x*, *t*) denote the concentration of a‘Response’ signal that originates from the apical ridge, corresponding to S1P.

The concentrations of the source *S* and the response *R*, are described by:

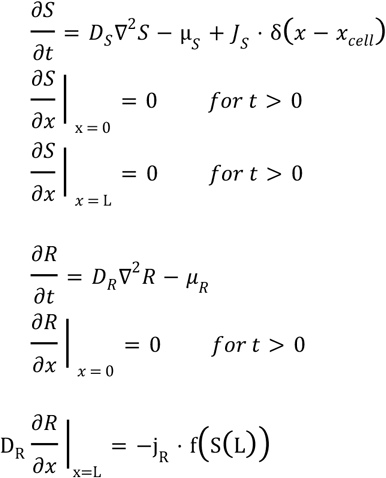

*S* is produced with rate *J_S_* at the position of the cell *x_cell_*, degrades with rate μ_*S*_, and diffuses with a diffusion coefficient *D_S_*. It has zero flux at the left and right boundaries. *R* is produced as a function of the amount of *S* on the right boundary, scaled by a production factor, -*j_R_*, diffuses with diffusion coefficient *D_R_* and degrades with rate μ_*R*_. It also has a zero-flux boundary condition on the left. L = 50 μm; X_cell_ (t = 0) = 5μm; D_S_, D_R_ = 10 μm^2^ s^-1^; μ_S_, μ_R_ = 0.3 s^-1^; J_S_, J_R_ = 0.3 mol s^-1^; γ = 4.5 x 10^-3^μm s^-1^; R_0_ = 5 x 10^-4^ mol

The cell migration rate *V_cell_* is a function of the amount of *R*present at the cell position, *R_cell_*. The migration rates of many cell types have been found to have a biphasic response to cell substrate adhesiveness. Maximum cell velocity takes place at intermediate levels of adhesiveness (Schwartz and Horwitz, 2006).

We model this dependence with the following velocity response function:

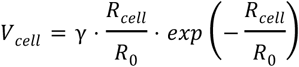

*R*_0_ is a characteristic concentration and *γ* is a constant that scales the velocity response. When R_cell_/R_0_ « 1 , it corresponds to a situation where the cell has weak contact with the substrate and insufficient traction, while *R_cell_*/*R*_0_ » 1 corresponds to the cell adhering very strongly to the substrate. Deeper analysis of the model will be included in a follow-up publication.

### Computer Simulations

Simulations were carried out in MATLAB R2018a by iteratively applying the bvp5c boundary value problem solver. We assume that the reaction-diffusion of signalling molecules *S* and *R* happens much faster than cell migration, such that the resulting distribution at each time step can be approximated by its steady state solution. For each time step, *R_cell_* is obtained through linear interpolation and used to calculate the cell position at the next step.

## ACKNOWLEDGEMENTS

Work in the TJC and TES labs was funded by an MOE Tier 3 grant (MOE2016-T3-1-005). Work in the FL lab was supported by funding from the National Science Foundation, IOS-1354457. SMP is supported by Senior Research Fellowships (1042589 and 1156693) from the National Health and Medical Research Council of Australia.

## SUPPLEMENTARY TABLE AND FIGURES

**Supplementary Table 1:** Variability in penetrance of fin blister phenotype in 7 different *stomp*^+/-^ incrosses.

|  | *stomp*<sup>+/-</sup> incross |  |  |  |  |  |  |
| --- | --- | --- | --- | --- | --- | --- | --- |
| Incross # | 1 | 2 | 3 | 4 | 5 | 6 | 7 |
| WT | 29 | 47 | 50 | 182 | 117 | 111 | 166 |
| Blisters | 8 | 7 | 7 | 16 | 4 | 5 | 34 |
| Total | 37 | 54 | 57 | 198 | 121 | 116 | 200 |
| Percent | 21.6 | 13.0 | 12.3 | 8.1 | 3.3 | 4.3 | 17.0 |

**Supplementary Figure S1:**
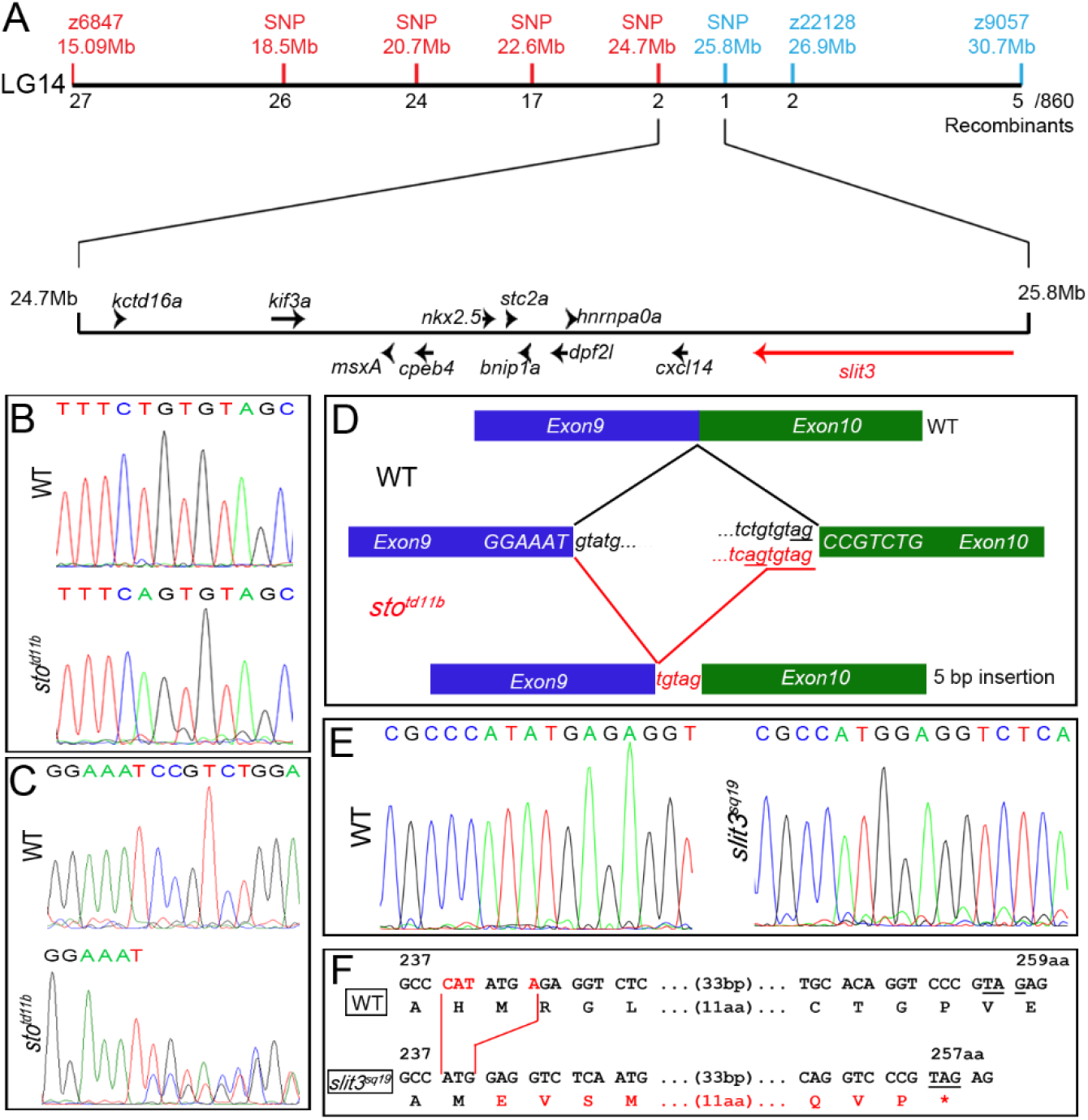
Mapping of *stomp* and TALEN mutagenesis of *slit3*. **A:** Linkage analysis using SSLP and SNP markers mapped *stomp* to LG14 with north and south markers represented in red and blue respectively with approximate chromosomal positions given. Genes and orientations within the interval are depicted below as arrows, including the causative gene, *slit3*, coloured in red. **B-D:** Sequence chromatograms of WT and *sto^td11b^* mutants DNA at the intron 9 - exon 10 splice junction of the *slit3* gene (B) and sequence of the corresponding region in the cDNA (C). The T>A generates a partially utilised cryptic splice site depicted in D leading to inclusion of 5 intronic nucleotides in the mutant cDNA (D – lower splicing in red). **E-F:** Characterisation of TALEN mutation of *slit3* at the DNA level showing sequence chromatograms of WT and the resulting *slit3^sq19^* allele (E) and the outcome at the cDNA level of WT (upper) and mutant allele (lower), showing deleted nucleotides in red and conceptual translation below (F).

**Supplementary Figure S2:**
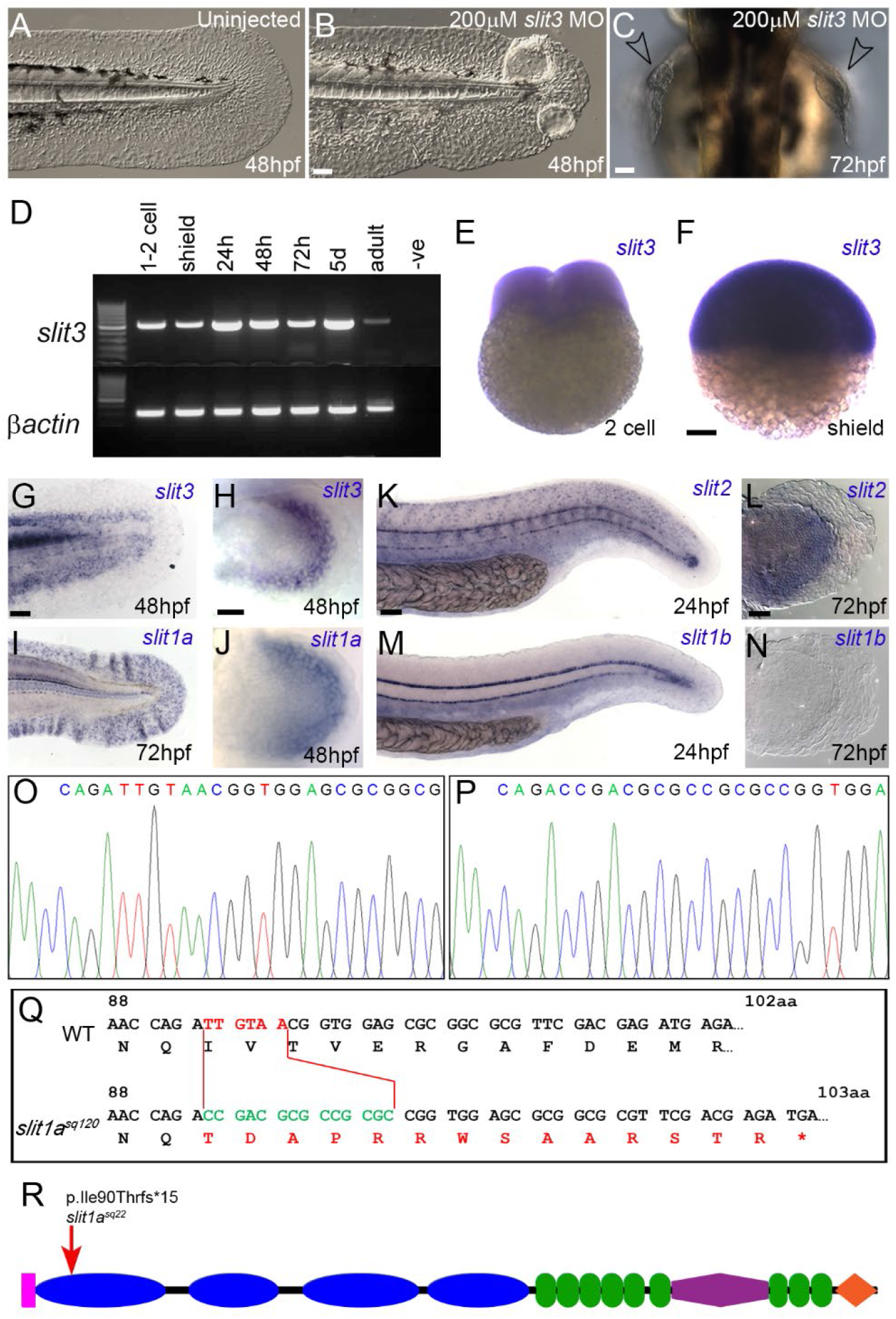
Expression of *slit* genes in the fins and mutagenesis of *slit1a*. **A-C:** Nomarski images of 48hpf tail (A, B) or 72hpf pectoral (C) fins imaged either laterally or dorsally. Larvae were either uninjected (A) or injected with 200μM *slit3* ATG translation blocking morpholino (B, C). Blisters are indicated in (C) with arrowheads. **D:** RT-PCR showing expression of *slit3* (upper panel) compared to *□actin* (lower panel) at all stages of zebrafish development including 1-2 cells stage. **E-N:** In situ hybridisation of 2 cell embryo (E), shield stage (F), tail fins (G, I, K, M) and pectoral fins (H, J, L, N) stained with probes against *slit3* (E-H), *slit1a* (I-J), *slit2* (K, L) and *slit1b* (M, N). Fin expression is limited to *slit3* and *slit1a*. **O-R:** CRISPR mutagenesis of *slit1a*.Sequencing chromatograms of PCR from gDNA of exon 3 of *slit1a* gene in WT (O) and mutants (P). The indel is presented in (Q) indicating the sequence of the WT (upper) and mutant (lower) alleles’ cDNA at the mutation site. Nucleotides deleted from the WT are shown in red and the inserted nucleotides given in green in the lower mutant allele. Corresponding translation presented below each with aberrant amino acids resulting from the frame shift presented in red below. The relative location of the mutation is presented on the protein domain schematic, with domains as per Figure 2A (R). Scale bars E-F: 100μm; Scale bars H-J, L-N: 20 μm. All other scale bars: 50μm.

**Supplementary Figure S3:**
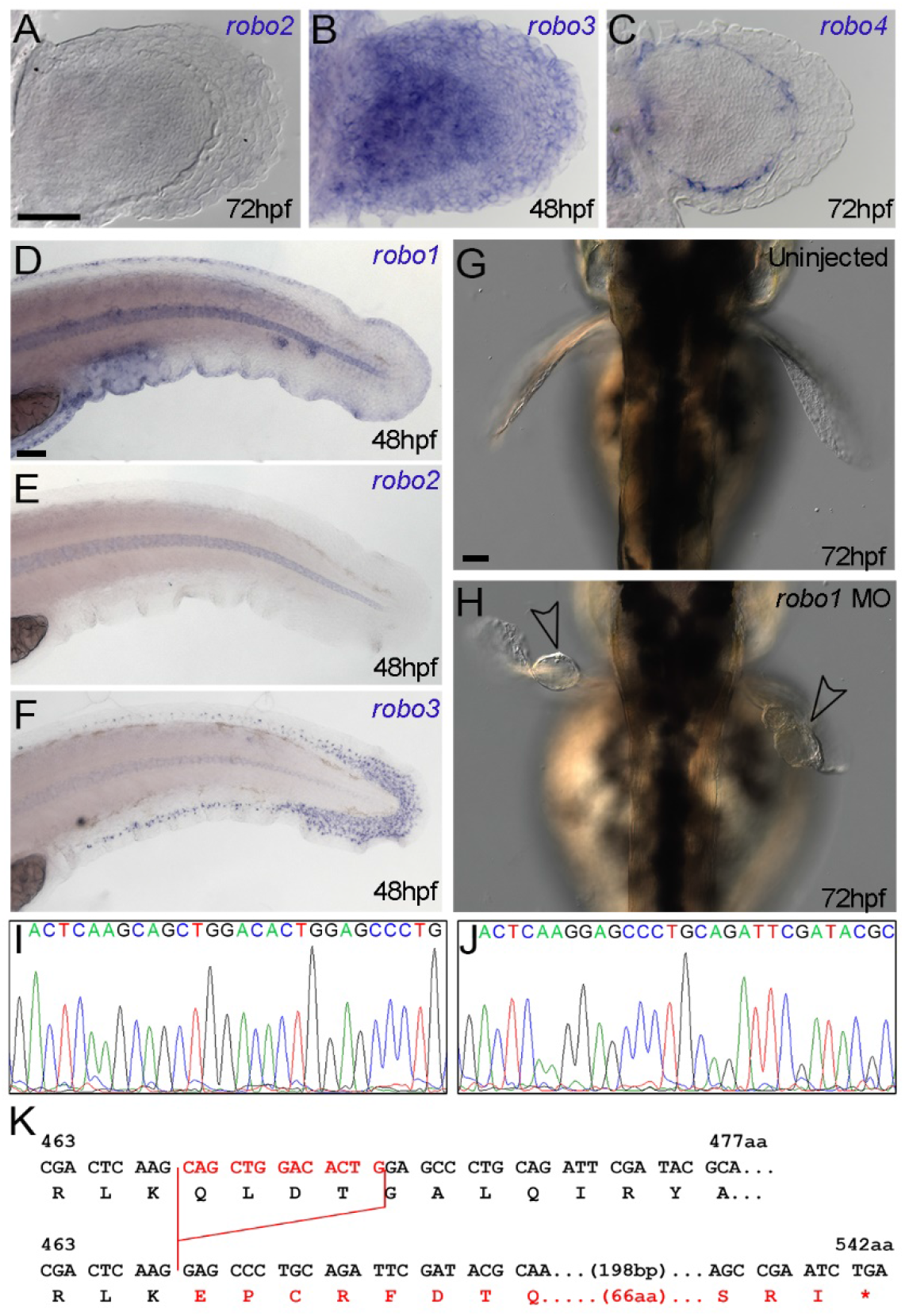
Expression of *robo* genes in the fins and strategies to disrupt Robo1 function. **A-F:** In situ hybridisation of the pectoral fins at 72hpf (A-C) and tail fins at 48hpf (D-F) stained using probes for *robo1* (D), robo2 (A, E), *robo3* (B, F) and *robo4* (C). While *robo1* expression persists at the fin apex, *robo2* is no longer expressed in the fins, *robo3* is broadly expressed in the pectoral fin, but has switched to mesenchyme expression in the tail, whilst *robo4* is expressed only in the pectoral fin vasculature. **G-H:** Dorsal Nomarski images of pectoral fins from 72hpf larvae uninjected (G) or injected with 500μM *robo1* ATG translation blocking morpholino (H). Blisters are indicated in (H) with arrowheads. **I-K:** TALEN mediated mutagenesis of exon 8 *robo1* gene. Sequencing chromatograms of TALEN targeted region in WT (I) and *robo1* allele (J). The deleted nucleotides are shown in red in (K) in the WT cDNA sequence (upper) with the resulting mutant *robo1* allele cDNA sequence shown below. Resulting translation is shown below the DNA sequences with the frameshifted mutant translation in red. Scale bars: 50μm.

**Supplementary Figure S4:**
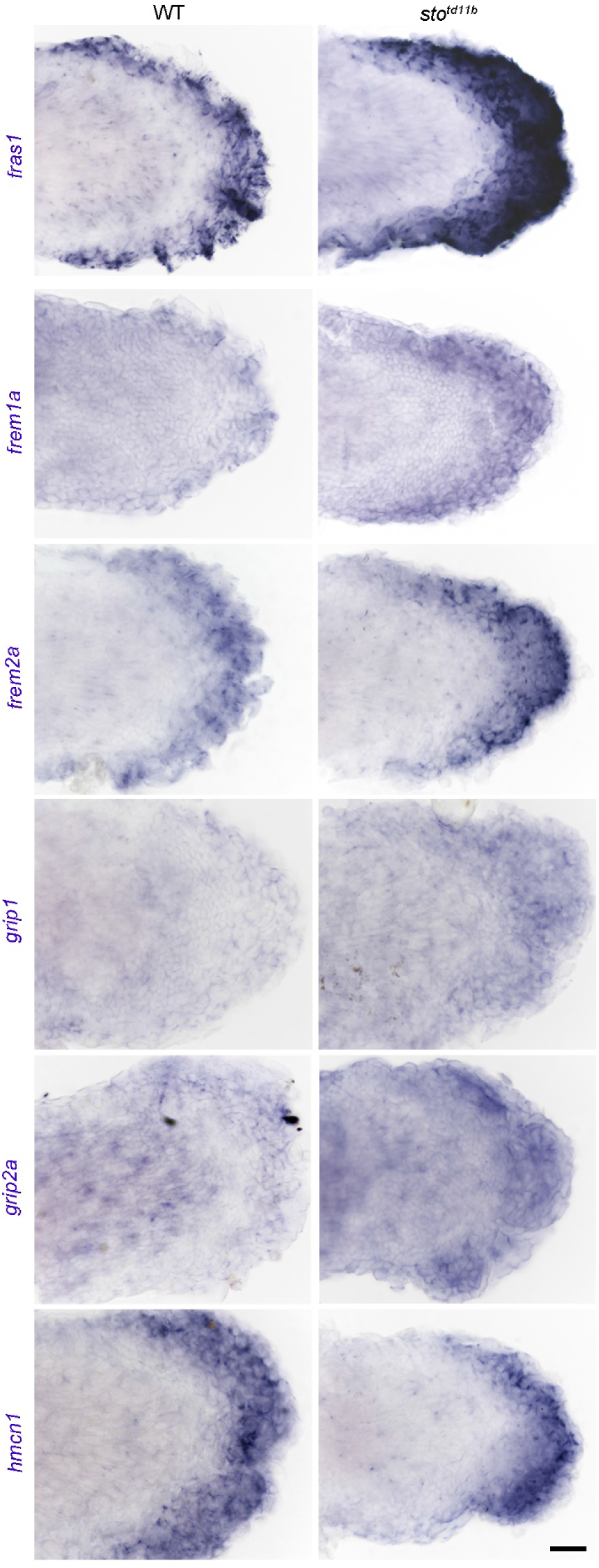
Expression of other blistering genes is not lost in *stomp*. In situ hybridisation of 72hpf pectoral fins of wild-type (left column) and *stomp* mutants (right column) stained with probes for (top row to bottom row) *fras1*, *frem1a*, *frem2a*, *grip1*, *grip2a* and *hmcn1*. Scale bar: 20μm.

**Supplementary Figure S5:**
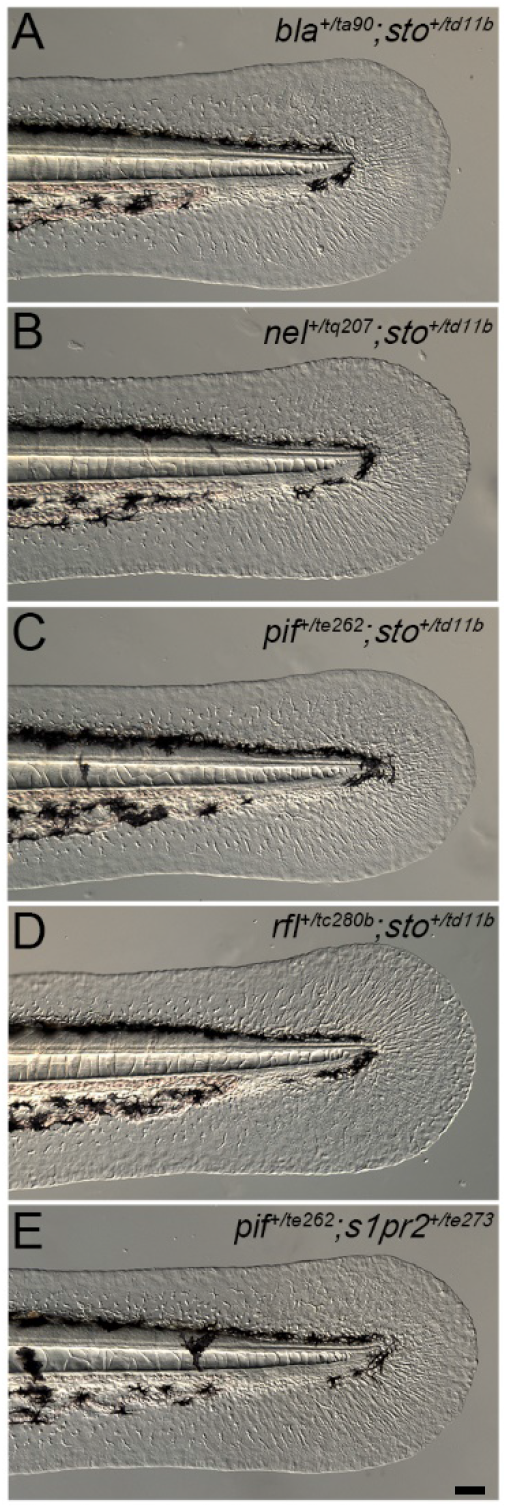
*stomp* complements Fraser complex genes. **A-E:** Lateral views of the tail region of embryos at 48hpf, which are trans-heterozygous for *sto*^+/*td11b*^ with other genes causing blistering, namely *bla*^+/*ta90*^ (*frem2a*; A), *nel*^+/*tq207*^ (*hemicentin1*; B), *pif*^+/*te262*^ (*fras1*; C), and *ffl*^+/*tc280b*^ (*frem1a*; D), as well as a trans-heterozygote for *pif*^+/*te262*^, with *miles apart* (*s1pr2*^+/*te273*^; E). In all cases trans-heterozygotes have WT fins, thus demonstrating that *stomp* is not allelic to, nor genetically interact with, these loci. In addition, *miles apart* does not genetically interact with *pinfin*. Scale bar: 50μm.

**Supplementary Figure S6:**
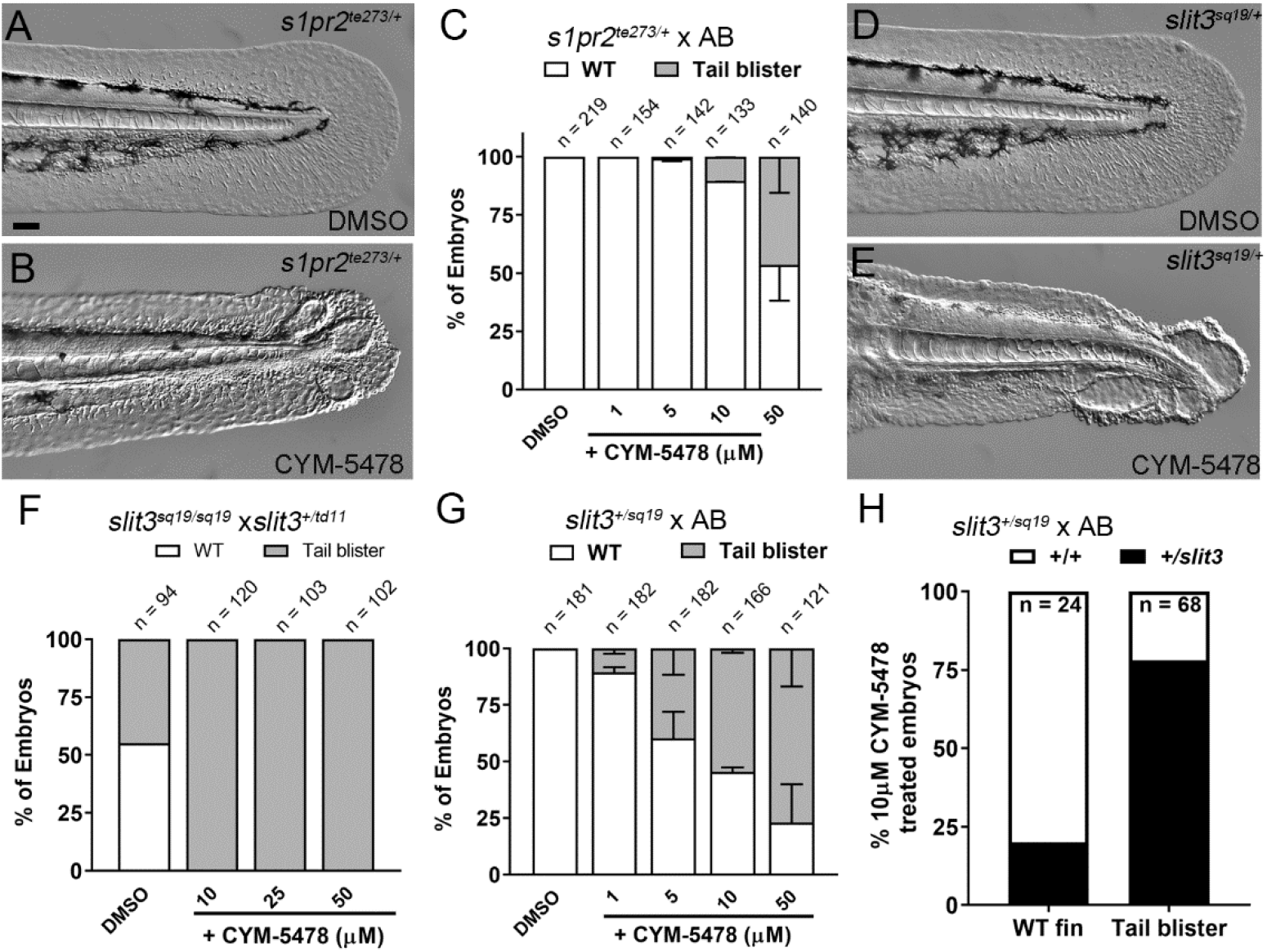
The S1PR2 modulator CYM-5478 synergises with reduced Slit3 and S1PR2 function. **A-C:** CYM-5478 synergises with *s1pr2* heterozygosity. Lateral Nomarski images of *s1pr2*^+/*te273*^ heterozygotes treated with DMSO (A) or 10μM CYM-5478 (B). CYM-5478 dose dependently induces blisters in *s1pr2*^+/*te273*^ heterozygotes (C). **D-E:**Lateral Nomarski images of the tail fins of *slit3*^+/*sq19*^ heterozygotes treated with 10μM CYM-5478 (E) or untreated (D). **F-H:** The proportion of larvae with tail blisters from a *slit3*^+/*sq19*^ outcross (G) or *slit3*^+/*sq19*^ crossed to a *slit3* mutant (F) and treated with given concentrations of CYM-5478. A dose dependant increase in blister frequency was observed, and those with blisters were predominantly *slit3* heterozygotes, whilst those unaffected by 10μM CYM-5478 were predominantly WT (H). Scale bar: 50μm.

**Supplementary Figure S7:**
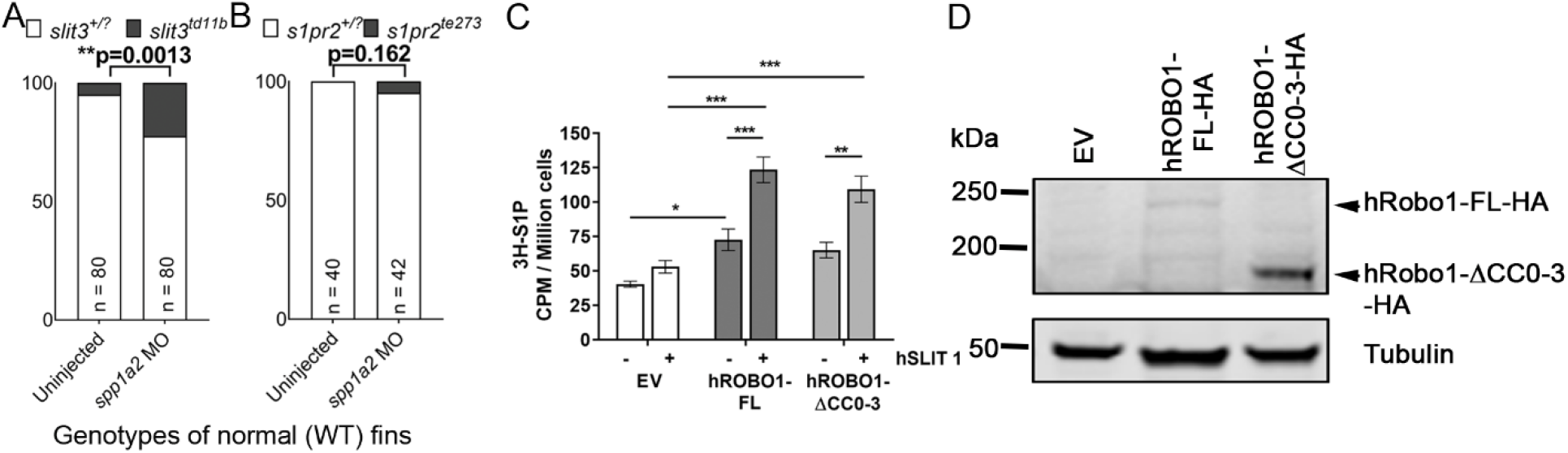
Slit-Robo signalling acts upstream of S1P signalling. **A-B:** Injection of *spp1a* and *spp2* morpholinos reduces the penetrance of fin blisters in *stomp* homozygotes, with significantly more *slit3^td11b^* mutants showing WT fins following morpholino injection compared to uninjected (A). Such a significant difference was not seen for *s1pr2^te273^* mutants (Chi-squared test). **C:** Intracellular S1P counts following transfection of HaCaT cells with empty vector (EV; white bars), full length Robo1 (dark grey bars) or truncated dominant negative Robo1 (light grey bars) and then metabolically labelling with 3H-sphingosine. Cells were stimulated with recombinant SLIT1 (+) or unstimulated (-). Radiolabelled intracellular S1P was measured by scintillation counting and corrected for cell number (*p<0.05; **p<0.005; ***p<0.001; ANOVA with Tukey’s post-test). **D:** Western Blot of the HaCaT cells used in (C) using an antibody against HA showing expression of the full length and truncated versions of ROBO1, compared to empty vector transfected cells and with β-Tubulin as a loading control.

**Supplementary Figure S8:**
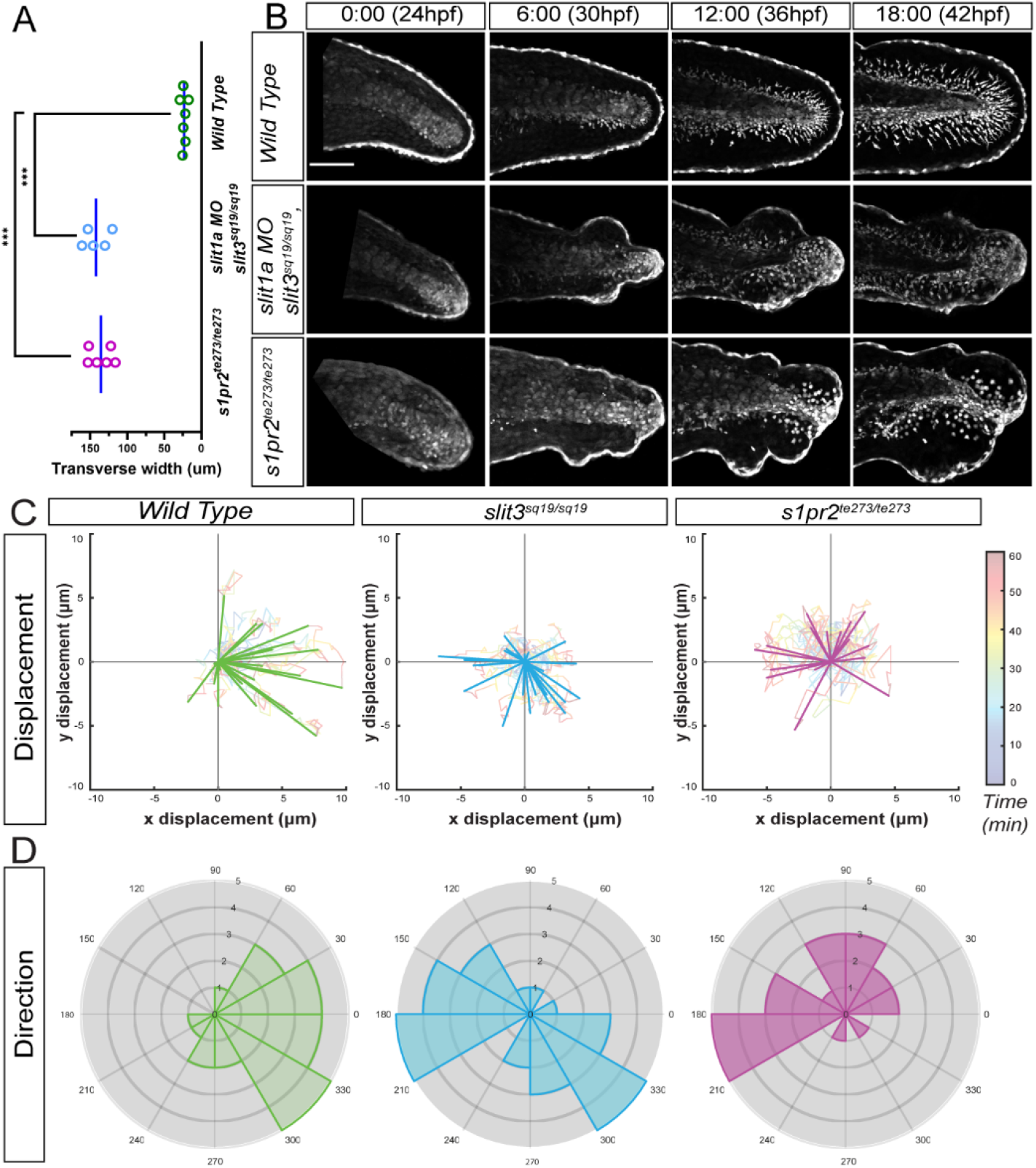
Similarities in cellular behaviour of mesenchymal cells in *slit3^sq19^* and *mil^te237^* mutants. **A:** Graph depicting the transverse width of the caudal fin edges of Wild Type, *slit3^sq19^* and *mil^te237^* mutants. **B:** Comparative Stills at every 6 hour intervals, from time-lapse confocal movies (Supplementary Movies 2, 3) of of WT (top), *slit3^sq19^* + *slit1a* MO (middle) and *s1pr2^te237^* (bottom), crossed to *sqet37Et*, labelling the mesenchymal cells in eGFP are shown. **C-D:** Magnitude (C) and direction histogram (D) of final cell displacement of cells from WT (left) *slit3*^-/-^ (centre) and *mil*^-/-^ (right) embryos over 60 minutes. The displacement measures in (C) are superimposed on the cell migration tracks. Mutants display reduced displacement and lack of directionality over a short-range.

**Supplementary Figure S9:**
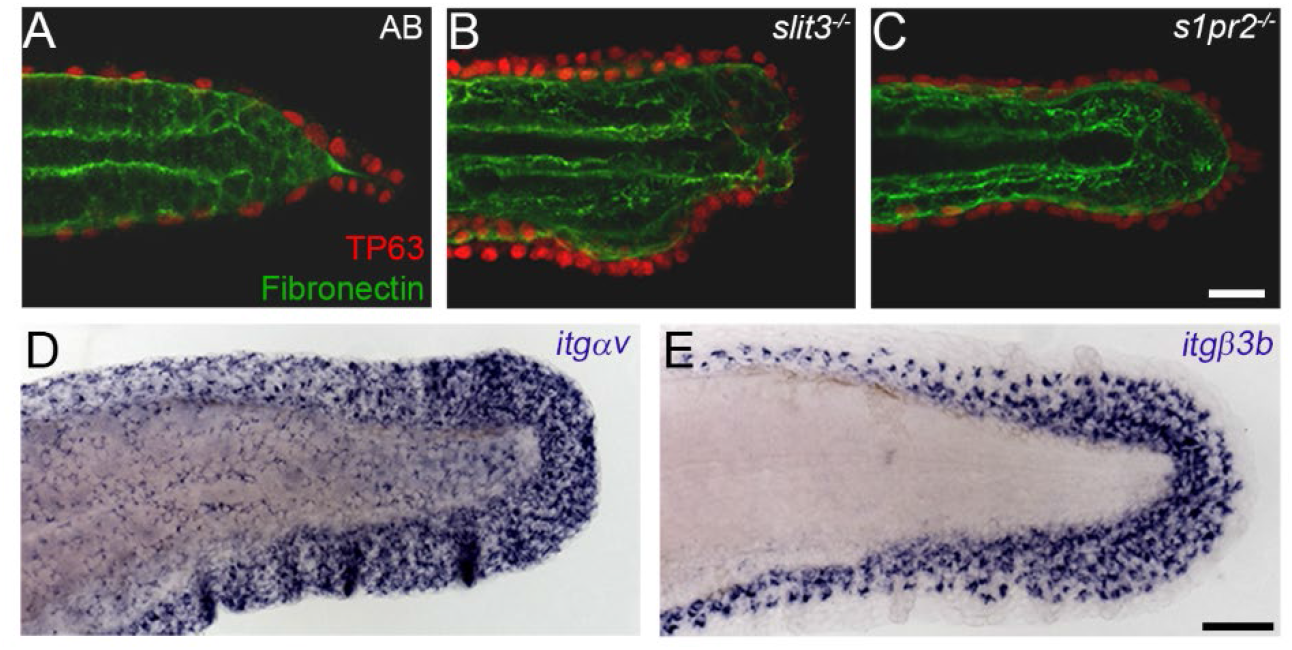
Fibronectin and its receptors are expressed in the fin interstitium and mesenchymal cells respectively. **A-C:** Confocal projections of dorsal views of 72hpf larvae immunostained for Fibronectin (green) and ΔNp63 (red) showing broad Fn staining under the epidermis of the fin and body WT at 72hpf. This staining does not appear reduced in *slit3*^-/-^ (B) or *s1pr2*^-/-^ (C) mutants. D-E: Lateral views of 36hpf tail fins stained by in-situ hybridisation for the fibronectin receptors *itgav* (D) and *itgβ3b* (E). Both are expressed in the fin mesenchymal cells, with *itgαv* additionally expressed in the epidermis as well. Scale bar A-C: 20μm; Scale bar **D-E**: 50μm.

## SUPPLEMENTARY MOVIES

**Supplementary Movie 1:**

**3D projections and rotation of the caudal fins of Wild Type, *slit3*^-/-^ and *s1pr2*^-/-^ at 40hpf.**

Confocal images were acquired at 20x magnification (0.5x Zoom) before being 3D projected in Imaris.

**Supplementary Movie 2:**

**Time lapse of Wild Type, *slit1a* MO + *slit3*^-/-^, and *s1pr2*^-/-^ embryos from 26hpf onwards.**

Confocal images were acquired at 20X magnification (0.5x Zoom) at every 10-minute interval for over 24 hours or until the blister collapsed (which ever was earlier). Time stamps indicate minutes and hours.

**Supplementary Movie 3:**

**Time lapse of individual mesenchyme cells traced in Wild Type, *slit1a* MO + *slit3*^-/-^, and *s1pr2*^-/-^ embryos from 30hpf onwards.**

Confocal images were acquired at 20X magnification (0.5x Zoom) at every 10-minute interval for over 24 hours or until the blister collapsed (which ever was earlier). Cell tracing was performed on such images, once every 5 frames (50 minutes) from 30hpf onwards. Time stamp indicates minutes. The embryos depicted here are the same as the ones shown in Supplementary Movie 2.

**Supplementary Movie 4:**

**Time lapse of mesenchyme cells in *Wild Type*, *slit3*^-/-^, and *s1pr2*^-/-^ embryos from 34hpf onwards.**

Images were acquired at 40X magnification (1x Zoom) at every 2-minute interval for 3 hours. Time stamps indicate minutes and hours.

